# ADAMTS7 promotes smooth muscle foam cell expansion in atherosclerosis

**DOI:** 10.1101/2024.02.26.582156

**Authors:** Allen Chung, Lauren E. Fries, Hyun-Kyung Chang, Huize Pan, Alexander C. Bashore, Karissa Shuck, Caio V. Matias, Juliana Gomez Pardo, Jordan S. Kesner, Hanying Yan, Mingyao Li, Robert C. Bauer

**Affiliations:** Cardiometabolic Genomics Program, Division of Cardiology, Department of Medicine, Columbia University, New York, NY, USA; Present address: Division of Cardiovascular Medicine, Department of Medicine, Vanderbilt University Medical Center, Nashville, TN, USA; Present address: Cardiovascular Research Institute, Icahn School of Medicine at Mount Sinai, New York, NY, USA; Department of Biostatistics, Epidemiology and Informatics, University of Pennsylvania Perelman School of Medicine, Philadelphia, PA, USA

## Abstract

Human genetic studies have repeatedly associated *ADAMTS7* with atherosclerotic cardiovascular disease. Subsequent investigations in mice demonstrated that ADAMTS7 is proatherogenic and induced in response to vascular injury. However, the cell-specific mechanisms governing ADAMTS7 proatherogenicity remain unclear. To determine which vascular cell types express *ADAMTS7*, we interrogated single-cell RNA sequencing of human carotid atherosclerosis and found *ADAMTS7* expression in smooth muscle cells (SMCs), endothelial cells (ECs), and fibroblasts. We subsequently created SMC- and EC-specific *Adamts7* conditional knockout and transgenic mice. Conditional knockout of *Adamts7* in either cell type does not reduce atherosclerosis, whereas transgenic induction in either cell type increases atherosclerosis. In SMC transgenic mice, this increase coincides with an expansion of lipid-laden SMC foam cells and a decrease in fibrous cap formation. RNA-sequencing in *Adamts7* overexpressing SMCs revealed an upregulation of lipid genes typically assigned to macrophages. Mechanistically, ADAMTS7 increases SMC oxLDL uptake through CD36, whose expression is upregulated by PU.1. ATAC-seq and motif analysis revealed increased chromatin accessibility at AP-1 enriched regions, consistent with AP-1 dependent remodeling of PU.1-regulated lipid-handling loci. In summary, ADAMTS7 promotes atherosclerosis by driving SMC foam cell formation through an AP-1/PU.1/CD36 regulatory axis.

## INTRODUCTION

Coronary artery disease (CAD) is the leading cause of death in the United States despite the widespread availability of multiple highly effective lipid lowering therapies (1). Thus, there remains a need for novel, non-lipid lowering therapeutic approaches for the treatment of CAD. Human genome-wide association studies (GWAS) have proven an effective, unbiased approach to discovering novel genomic loci associated with CAD, potentially uncovering therapeutic targets. GWAS have repeatedly identified the 15q25 locus as a region of interest for CAD (2–5). This locus harbors the gene <u>A</u> <u>d</u>isintegrin <u>a</u>nd <u>m</u>etalloproteinase with thrombo<u>s</u>pondin motifs <u>7</u> (*ADAMTS7*), which encodes a secreted matrix metalloproteinase previously implicated in osteoarthritis (6). After its identification by GWAS, subsequent mouse studies demonstrated that ADAMTS7 is proatherogenic, as mice with whole-body *Adamts7* knockout had reduced atherosclerosis (7). Notably, this reduction in atherosclerosis occurred without any changes in plasma lipoprotein levels, suggesting that ADAMTS7 has a lipid-independent role in atherogenesis (8). Further mouse models of *Adamts7* have confirmed this proatherogenicity while demonstrating that ADAMTS7 catalytic activity is required for this effect (9, 10). Collectively, these findings identify ADAMTS7 as an attractive lipid-independent therapeutic target for CAD, although the molecular mechanisms linking ADAMTS7 to atherogenesis remain incompletely understood.

There has recently been an increased appreciation for the role of smooth muscle cells (SMCs) in atherosclerosis (11). Although historically viewed primarily as contributors to fibrous cap formation, single-cell RNA sequencing (scRNA-seq) studies in mice and humans have revealed that SMCs are highly plastic and capable of undergoing phenotypic modulation, acquiring features of multiple cell types, including macrophage-like characteristics (12, 13). Consistent with this plasticity, studies in the *Apoe*^-/-^model of atherosclerosis demonstrated that most lipid-laden aortic foam cells are not macrophages but cells of SMC origin, highlighting that SMC foam cells may play a more significant role in atherogenesis than previously understood (14). Mechanistic studies of ADAMTS7 have largely focused on its effects on SMC function, as both *Adamts7* knockout and overexpression alter neointima formation following vascular injury and modulate SMC migration in ex vivo assays (7, 15). As ADAMTS7 is a secreted protein, it may influence multiple vascular cell types through both cell-autonomous and paracrine mechanisms (16). *Adamts7* is not constitutively expressed in vascular cells, but rather, its expression is induced in response to vascular injury and cytokine signaling (7, 9). Low basal expression of *Adamts7* in vivo, together with its transient induction, has hindered efforts to identify the cell types that express *Adamts7* and to define its function during atherosclerosis (7). Overall, ADAMTS7 has consistently been linked to SMC function, yet the mechanistic basis by which this association promotes atherosclerosis remains unknown (8).

In this study, we identified *ADAMTS7* expression across multiple human vascular cell types, including SMCs and endothelial cells (ECs). We additionally generated several novel tissue-specific *Adamts7* mouse models, enabling mechanistic interrogation of ADAMTS7 under conditions of sustained induction or targeted deletion. Using these models, we demonstrate that *Adamts7* expression in either SMCs or ECs is individually sufficient to increase atherosclerosis and promote SMC foam cell formation, whereas deletion in either cell type alone does not significantly affect lesion burden. Mechanistically, we found that ADAMTS7 alters the chromatin landscape of SMCs, inducing expression of the myeloid transcription factor (TF) PU.1, increasing *Cd36* expression, and enhancing SMC oxidized low-density lipoprotein (oxLDL) uptake. Together, these findings demonstrate that ADAMTS7 drives SMC foam cell expansion and, ultimately, atherosclerosis.

## RESULTS

### *ADAMTS7* is induced within multiple vascular cell types during lesion formation

Prior literature indicates that *Adamts7* expression is induced within the vasculature and promotes SMC migration while inhibiting re-endothelialization (7, 17). ADAMTS7 is secreted (18), and as such, the cell type that responds to it does not necessarily have to produce it. Thus, to determine the cellular origins of ADAMTS7, we interrogated the largest human atherosclerotic carotid artery scRNA-seq dataset generated to date (19) and found that multiple vascular cell types express *ADAMTS7* in mature plaques. The highest *ADAMTS7* expression was present in ECs, whereas SMCs, fibroblasts, and mast cells displayed *ADAMTS7* expression at a lower level (Figure 1A). Interestingly, the highest level of *ADAMTS7* expression occurred in ECs annotated as inflammatory (19), whereas fibrotic ECs did not display as high a level of expression.

**Figure 1.**
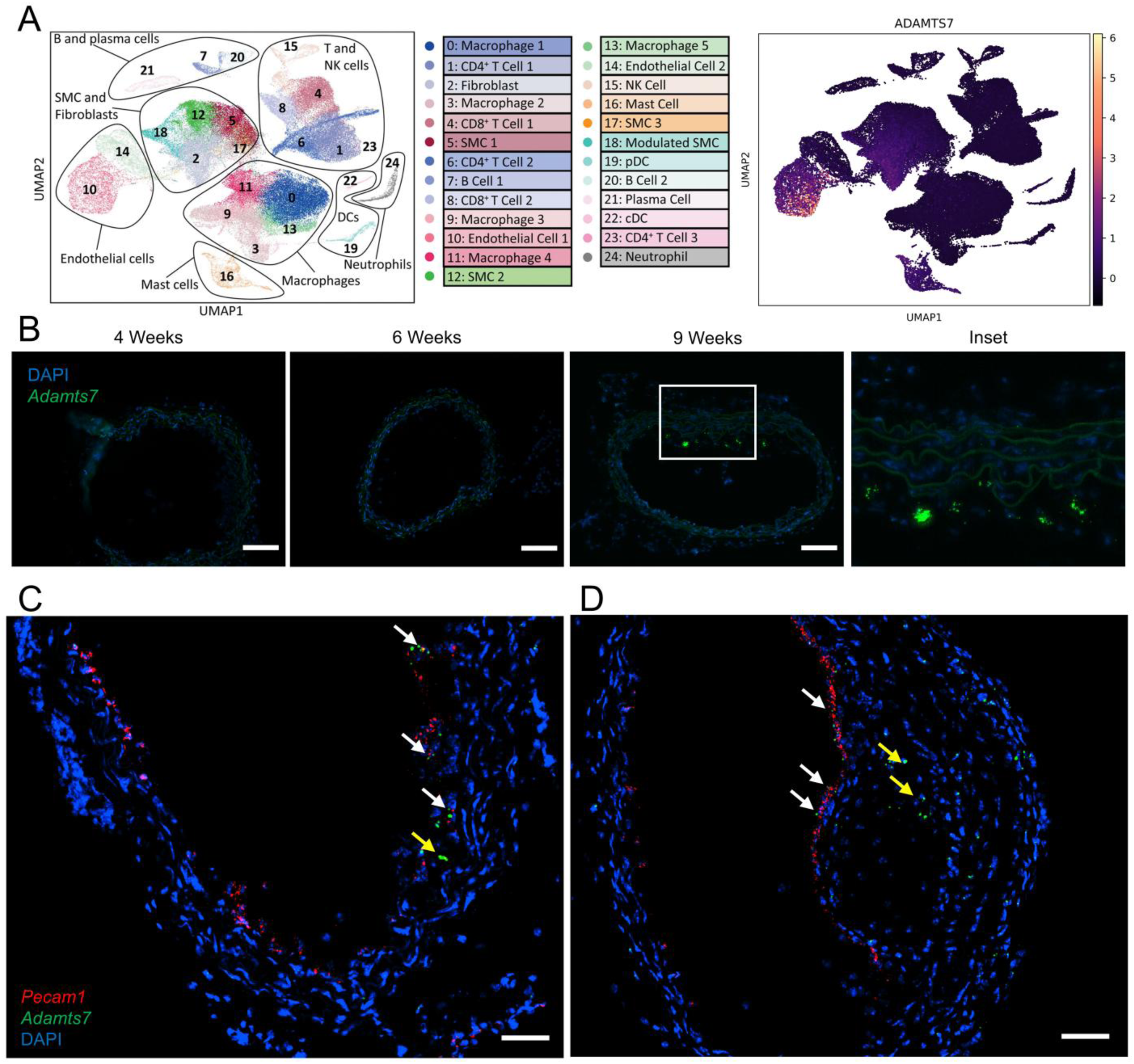
Identifying endogenous ADAMTS7 expression. (A) Single Cell RNA sequencing of human carotid atherosclerosis and cell clustering identities. Cluster identities are reproduced with permission from Bashore AC et al., Arterioscler Thromb Vasc Biol. 2024;44:930–945. (B) RNAscope of the BCA in mice after WTD feeding. Nuclei are stained with DAPI. Scale bar = 100 µm. RNAscope of the BCA of *Ldlr^KO^* mice after seven (C) and ten (D) weeks of WTD feeding and probed against *Adamts7* and *Pecam1*. White arrows highlight EC expression of *Adamts7*. Yellow arrows indicate non-EC expression of *Adamts7*. Scale bar = 50 µm.

We next sought to confirm this observation in mouse models of atherosclerosis. Previous work in *Apoe*^-/-^ mice has shown that *Adamts7* expression is induced after four weeks of western-type diet (WTD) feeding but absent by terminal time points (7). To determine when *Adamts7* is expressed during atherosclerosis, we performed RNAscope within the brachiocephalic artery (BCA) of hyperlipidemic Low-Density Lipoprotein Receptor knockout (*Ldlr*^KO^) mice at four, six, and nine weeks of WTD. *Adamts7* expression appears during early atheroma formation and is especially prominent at nine weeks when established lesions are present (Figure 1B). Across multiple experiments, *Adamts7* expression was localized to the developing atherosclerotic neointima and to multiple other locations in the BCA, consistent with its expression in SMCs and other cells (Figure 1, B-D). To confirm EC expression as seen in the human scRNA-seq data, we performed RNAscope in the BCA of *Ldlr*^KO^ mice after seven and ten weeks of WTD (Figure 1, C and D). At the early lesion time point of seven weeks, we detected colocalization of the *Adamts7* transcript with *Pecam1*, an EC marker, while at the ten-week time point, we observed *Adamts7* expression in both ECs and non-ECs. These data support the notion that multiple vascular cell types produce ADAMTS7 during atherosclerosis in mice and humans.

### SMC or EC specific knockout of *Adamts7* does not reduce atherosclerosis

Given *Adamts7* expression in multiple vascular cell types, we next asked if the knockout of *Adamts7* in any single vascular cell type could reduce atherosclerosis. We generated a conditional knockout model of *Adamts7* where exons five and six are floxed (Supplemental Figure 1A). Given the strong prior literature on ADAMTS7 regulation of SMC function, we crossed this mouse to the *Myh11*-CreER^T2^ and *Ldlr*^KO^ to generate a hyperlipidemic SMC conditional knockout of *Adamts7* (*Adamts7*^fl/fl^; *Ldlr*^KO^; *Myh11*-CreER^T2^, referred to as *Adamts7*^SMCKO^ *Ldlr*^KO^) and subsequently induced knockout through intraperitoneal tamoxifen injections (Figure 2A). Although *Adamts7* expression is typically low in uninjured vessels, SMC explants from *Adamts7*^SMCKO^ *Ldlr*^KO^ mice exhibited an approximately 50% reduction in *Adamts7* expression compared with wild-type controls *Adamts7*^SMCWT^ *Ldlr*^KO^ (Supplemental Figure 1B). Following 16 weeks of WTD feeding, we observed no changes in body weight or total cholesterol (Supplemental Figure 1, D and E). Surprisingly, *Adamts7*^SMCKO^ *Ldlr*^KO^ did not alter the lesion area in either en face Oil Red O (ORO) analysis (Figure 2B) or aortic root lesion area (Figure 2C). Because *ADAMTS7* expression is also prominent in ECs based on human scRNA-seq data (Figure 1A), we next generated an *Adamts7* EC-specific conditional knockout by crossing the hyperlipidemic conditional knockout mouse to the *Cdh5*-CreER^T2^ (*Adamts7*^fl/fl^; *Ldlr*^KO^; *Cdh5*-CreER^T2^, referred to as *Adamts7*^ECKO^ *Ldlr*^KO^). To confirm EC-specific knockout, we isolated CD31^+^ ECs from aortas and observed markedly reduced *Adamts7* expression in the CD31^+^ fraction from *Adamts7*^ECKO^ *Ldlr*^KO^ mice compared with wild-types (*Adamts7*^ECWT^ *Ldlr*^KO^), whereas no difference was observed in the CD31^−^ (non-EC) fraction (Supplemental Figure 1C). After 16 weeks of WTD feeding, we assessed atherosclerosis in *Adamts7*^ECKO^ *Ldlr*^KO^ compared to *Adamts7*^ECWT^ *Ldlr*^KO^ (Figure 2D). Again, we observed no changes in body weight or total cholesterol levels (Supplemental Figure 1, F and G). Similar to *Adamts7*^SMCKO^ *Ldlr*^KO^ mice, EC knockout of *Adamts7* also did not demonstrate any changes in the atherosclerotic burden (Figure 2, E and F). These data strongly suggest that individual knockout of *Adamts7* in ECs or SMCs alone is not adequate to alter atherogenesis in mice.

**Figure 2.**
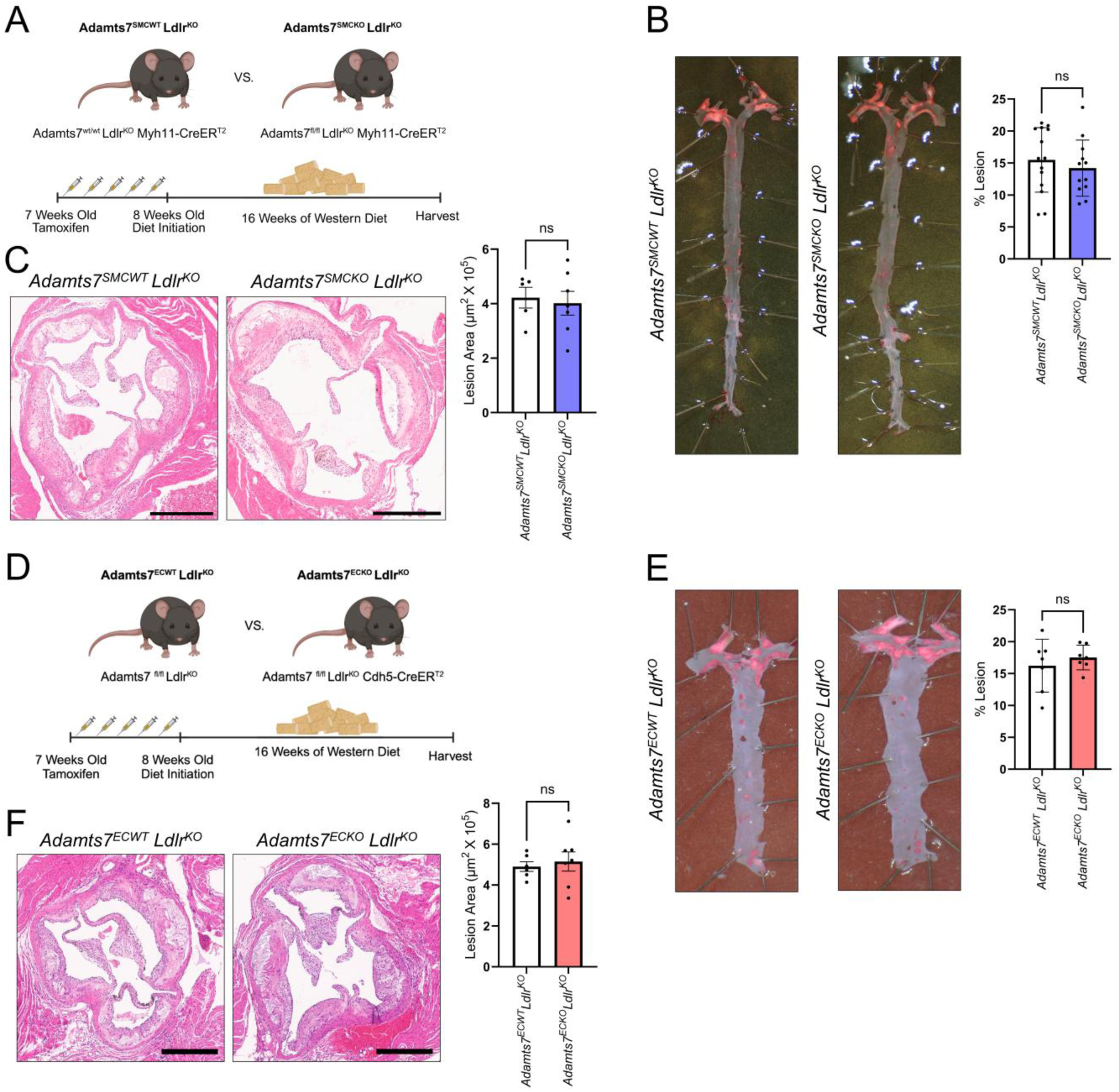
SMC or EC knockout of *Adamts7* does not affect atherosclerosis. (A) Schematic outlining experimental design and mouse comparisons for SMC knockout of *Adamts7*. Created with BioRender.com (B) Representative ORO staining of en face aortas and enumeration. *n =* 12 - 14. (C) Representative images of hematoxylin and eosin (H&E) stained aortic root sections, quantification of plaque areas *n =* 5 - 7. Scale bar = 500 µm. (D) Schematic outlining experimental design and mouse comparisons for EC knockout of *Adamts7*. Created with BioRender.com (E) Representative ORO staining of en face aortas and enumeration. *n =* 7. (F) Representative images of H&E stained aortic root sections, quantification of plaque areas *n =* 6 - 7. Scale bar = 500 µm. Statistics were analyzed using a 2-tailed Student’s t-test.

### Transgenic overexpression of *Adamts7* in mouse SMCs or ECs exacerbates atherosclerosis

Given that knockout of *Adamts7* in either SMCs or ECs alone does not appear to reduce atherogenesis in our model, we next asked whether sustained induction of *Adamts7* expression in a single cell type can increase atherogenesis. As *ADAMTS7* is induced in multiple vascular cell types in human atherosclerosis (Figure 1A), we generated a transgenic mouse model with conditional *Adamts7* overexpression to mimic its induction and promote its function in multiple cell types. We inserted murine *Adamts7* into the Rosa26 locus with a lox-stop-lox (LSL) cassette preceding the gene, allowing for tissue type-specific overexpression of *Adamts7*. The transgenic overexpression model was subsequently bred to the *Ldlr*^KO^ background as well as to either the *Myh11*-CreER^T2^ (*Adamts7*^SMCTG^) or the *Cdh5*-CreER^T2^ (*Adamts7*^ECTG^) to allow for SMC (Figure 3A) or EC (Figure 3G) specific overexpression of *Adamts7,* respectively. We verified *Adamts7* overexpression in the SMC transgenic model via qPCR (Supplemental Figure 2A) and western blot (Supplemental Figure 2B) from isolated aortic RNA and protein. We note that *Adamts7*^SMCTG^ *Ldlr*^KO^ exhibited a large-fold change in *Adamts7* RNA, but this is partly due to the near complete absence of baseline expression of *Adamts7*. Primary SMC explants from *Adamts7*^SMCTG^ displayed increased migration (Supplemental Figure 2C), consistent with prior findings (7). Validation of *Adamts7*^ECTG^ was achieved by examining *Adamts7* expression in isolated CD31^+^ vascular ECs as well as mouse lung ECs (Supplemental Figure 2D); *Adamts7* expression was elevated in ECs, whereas CD31^-^ cells showed no difference in *Adamts7* expression.

**Figure 3.**
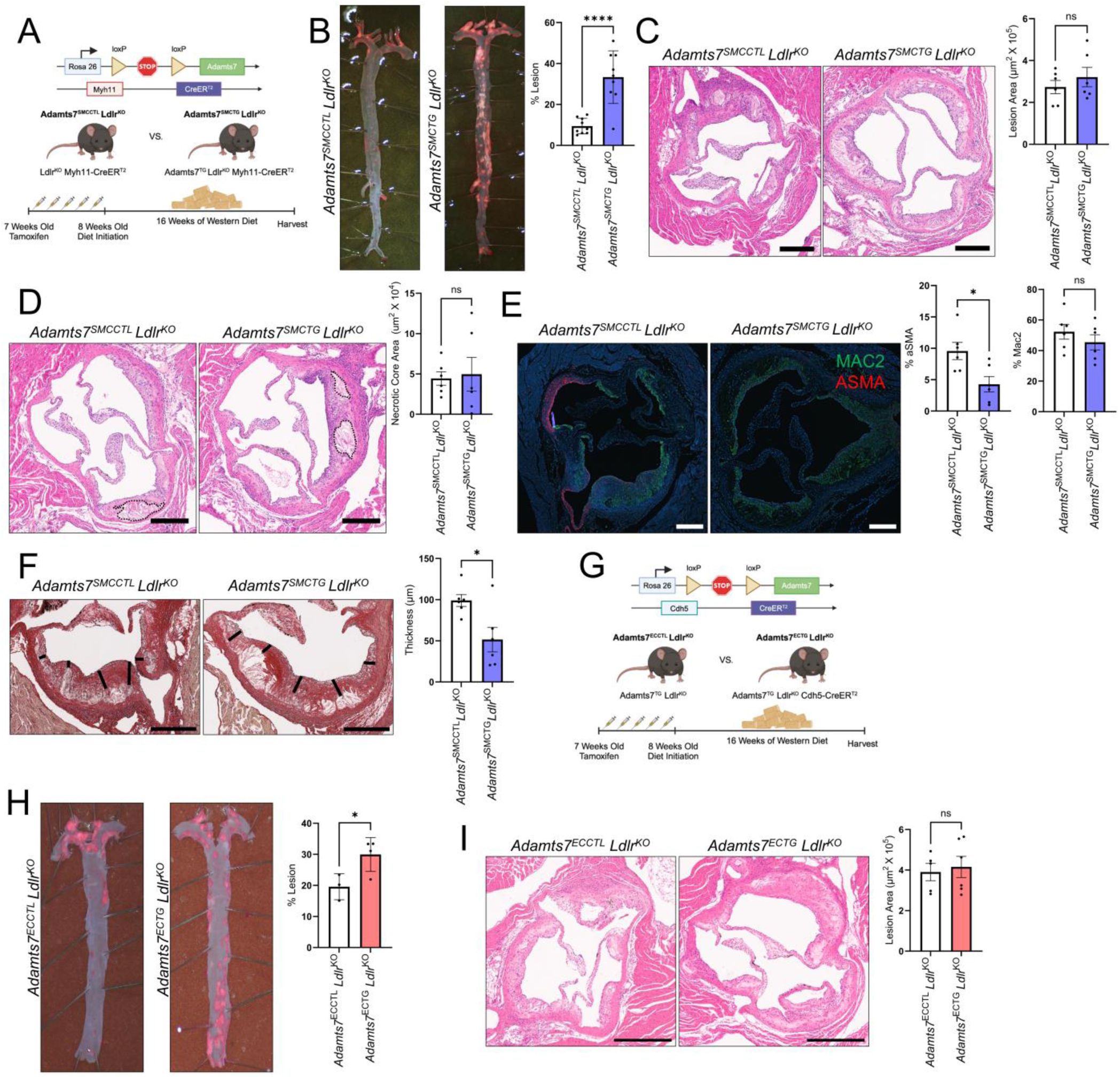
SMC and EC transgenic *Adamts7* increases aortic atherosclerosis. (A) Schematic outlining experimental design and the generation of the SMC transgenic ADAMTS7 mouse. Created with BioRender.com (B) ORO staining of the en face aorta and quantification of lesion area. *n =* 9. (C) Representative images of H&E stained aortic root sections, quantification of plaque areas *n =* 6. Scale bar = 500 µm. (D) Representative images of H&E stained aortic root sections with necrotic core outlined in dotted line *n =* 6 mice. Scale bar = 300 µm. (E) Representative images of aortic root sections stained against α-SMA (Cy3 - red), MAC2 (Alexa Fluor 488 - green), and cell nuclei (DAPI - blue) with their subsequent quantification relative to lesion area *n =* 6. Scale bar = 300 µm. (F) Representative picrosirius red staining of aortic root lesions with bars indicating fibrous cap thickness and quantification of fibrous cap thickness *n =* 6. Scale bar = 300 µm. (G) Schematic outlining experimental design and the generation of the EC transgenic ADAMTS7 mouse. Created with BioRender.com (H) ORO staining of the en face aorta and quantification of lesion area. *n =* 3 - 4. (I) Representative images of H&E stained aortic root sections, quantification of plaque areas *n =* 5 - 6. Scale bar = 500 µm. \*\*\*\**P*<0.0001, \*\*\**P*<0.001, ** *P*<0.01, \**P*<0.05. Statistics were analyzed using a 2-tailed Student’s t-test.

To investigate the role of sustained cellular *Adamts7* in atherosclerosis development, *Adamts7* overexpression was induced in all animal models via tamoxifen injection, and mice were fed a WTD for 16 weeks. There were no plasma cholesterol or body weight changes in any animal models (Supplemental Figure 3, E-H). *Adamts7*^SMCTG^ *Ldlr*^KO^ mice had a stark 3.5-fold increase in atherosclerosis as shown by en face ORO staining as compared to control *Adamts7*^SMCCTL^ *Ldlr*^KO^ mice (Figure 3B), but no change in the aortic root lesion area as assessed by H&E staining (Figure 3C), consistent with findings in other *Adamts7* mouse models (9). Given the dramatic increase in aortic atherosclerosis, we asked whether shorter durations of WTD feeding would confer an atherosclerosis phenotype. Indeed, we found increased atherosclerosis at three, six, and nine weeks of WTD feeding (Supplemental Figure 3A).

During atherosclerosis, SMCs play an essential role in plaque stabilization (20). As such, we further characterized the aortic root plaque morphology in *Adamts7*^SMCTG^ *Ldlr*^KO^ mice. We found no changes in necrotic core area (Figure 3D) or macrophage content via MAC2 staining (Figure 3E). However, *Adamts7*^SMCTG^ *Ldlr*^KO^ lesions had reduced SMC content as determined by α-SMA staining (Figure 3E), especially in the fibrous cap region. The reduction in α-SMA also coincided with a 48% reduction in fibrous cap thickness as assessed through picrosirius red staining (Figure 3F). The lack of change in MAC2 staining and the reduction in cap thickness indicate that increased ADAMTS7 reduces indices associated with human plaque stability in these mouse models. Finally, in tandem, we performed a similar atherosclerosis study on the *Adamts7*^ECTG^ *Ldlr*^KO^ mice and observed a 1.5-fold increase in aortic atherosclerosis in *Adamts7*^ECTG^ *Ldlr*^KO^ mice as compared to controls (Figure 3H). We also observed the same lack of changes in aortic root lesion area between groups (Figure 3I).

### *Adamts7* promotes SMC foam cell formation

The significant increase in ORO staining of the en face aorta indicates an increased accumulation of neutral lipids within vascular cells, presumably due to an expansion in foam cell formation. Thus, we next sought to assess aortic foam cell content in our *Adamts7* transgenic mice using a neutral lipid dye, LipidTOX. After excluding debris, doublets, and dead cells (Supplemental Figure 4A), we observed that the *Adamts7*^SMCTG^ *Ldlr*^KO^ had a threefold increase in foam cell content, mirroring the rise in aortic atherosclerosis (Figure 4A). Notably, over 80% of these foam cells lacked CD11B and CD64, indicating that non-leukocytes are accumulating lipids and becoming foamy (Figure 4B) (21). Given that increased ORO staining of the aorta was evident even at early time points, we next examined foam cell composition after 3 weeks of WTD. Despite the short feeding period, *Adamts7*^SMCTG^ *Ldlr*^KO^ mice already exhibited a significant increase in foam cell formation, driven primarily by an expansion of SMC-derived foam cells as identified by CD200 expression (Supplemental Figure 3B).

**Figure 4.**
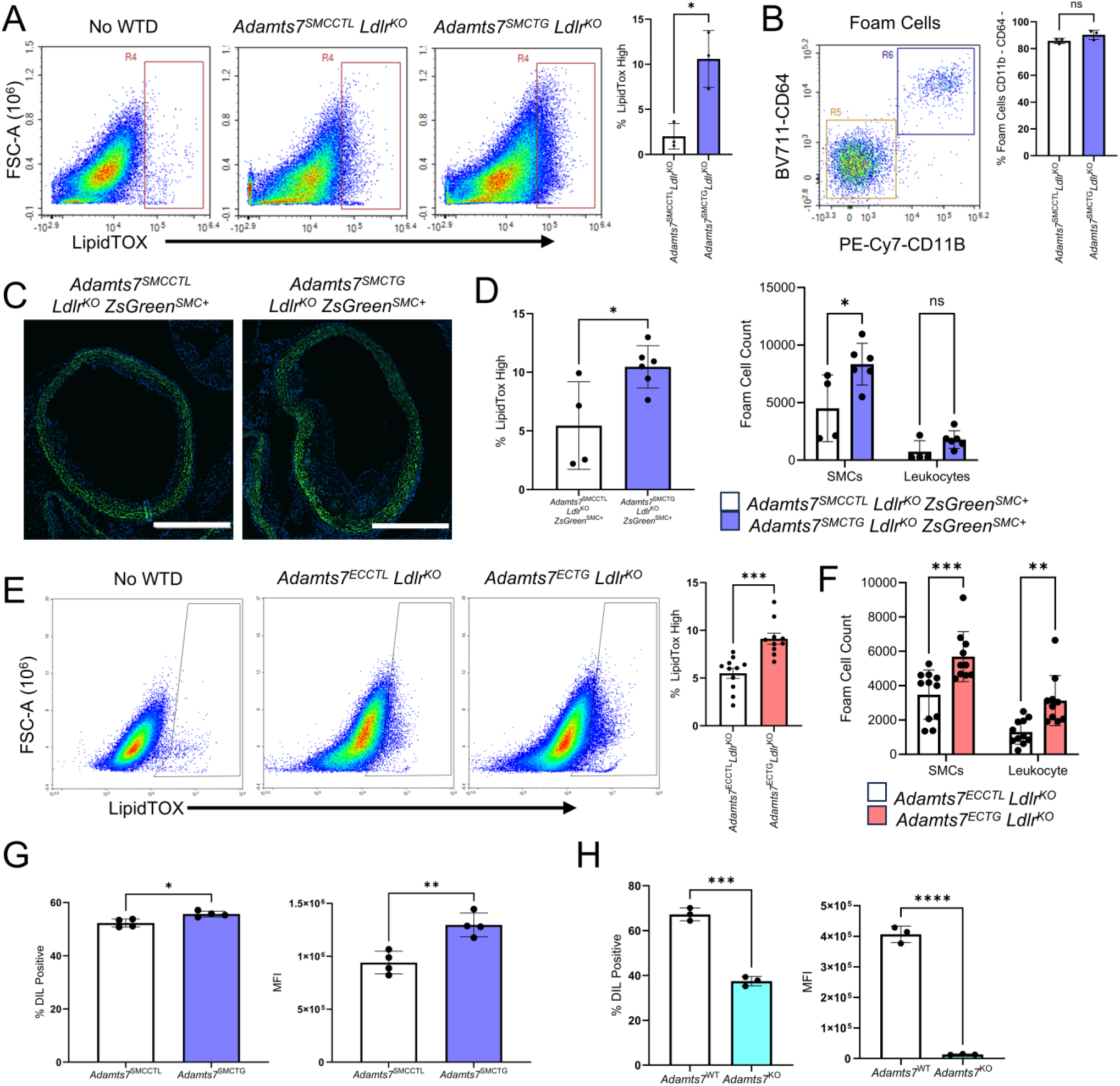
ADAMTS7 promotes foam cell expansion. (A) Flow cytometry-based quantification of foam cells through LipidTOX staining of the aorta. A normolipidemic aorta was used to establish the LipidTOX high gate. *n =* 3 (B) Breakdown of foam cells stained positively for CD11B and CD64 *n =* 3. (C) Confirmation of efficient ZsGreen labeling of SMCs with nuclei counterstained with DAPI. Scale bar = 500 µm. (D) Foam cell analysis with the SMC lineage tracer ZsGreen and leukocyte marker CD45 after 12 weeks of WTD. Counts were normalized to 100,000 live cells. *n =* 4 - 6 mice. Statistics were analyzed by two-way ANOVA with Sidak’s multiple comparison test. (E) Flow cytometry-based quantification of foam cells through LipidTOX staining of the aorta. A normolipidemic aorta was used to establish the LipidTOX high gate. *n =* 3 - 4. (F) Normalized foam cell counts out of 100,000 live cells after 16 weeks of WTD. Leukocytes were identified as CD45^+^. SMCs were identified as CD31^-^ CD45^-^ CD200^+^. *n =* 10 - 11 mice. Statistics were analyzed by two-way ANOVA with Sidak’s multiple comparison test. (G) In vitro foam cell analysis with primary cells from transgenic mice treated with 10 µg/mL DiI-oxLDL for 24 hours, *n =* 4. (H) In vitro foam cell analysis with explanted primary SMCs from whole body *Adamts7* knockout mice treated with 10 µg/mL DiI-oxLDL for 24 hours, *n =* 3. \*\*\*\**P*<0.0001, \*\*\**P*<0.001, ** *P*<0.01, \**P*<0.05 Between sample comparisons were analyzed by a 2-tailed Student’s t-test.

To further confirm that the increase in foam cells was predominantly of SMC origin, the *Adamts7*^SMCCTL^ *Ldlr*^KO^ and *Adamts7*^SMCTG^ *Ldlr*^KO^ mouse model was subsequently bred to an LSL-ZsGreen reporter (*Adamts7*^SMCCTL^ *Ldlr*^KO^ ZsGreen^SMC+^ and *Adamts7*^SMCTG^ *Ldlr*^KO^ ZsGreen^SMC+^) to allow for the lineage tracing of SMCs and their derived cells (12). Efficient ZsGreen labeling of SMCs was achieved through microscopy (Figure 4C). After 12 weeks of WTD feeding, ZsGreen status and concurrent staining with CD45, revealed that over 65% of lipid-laden foam cells were ZsGreen^+^ and CD45^-^ (Supplemental Figure 4B), indicating again that *Adamts7* was predominately increasing SMC foam cell formation (Figure 4D). We repeated the foam cell analysis within *Adamts7*^ECCTL^ *Ldlr*^KO^ and *Adamts7*^ECTG^ *Ldlr*^KO^ and observed that the *Adamts7*^ECTG^ *Ldlr*^KO^ mice contained an increase in foam cell content as well (Figure 4E). Further assessment of these foam cells within the aorta revealed that the majority were CD45^-^ and CD31^-^ while being CD200^+^ (21), indicating again that SMCs are also becoming foamy in this model (Figure 4F).

To test if ADAMTS7 promotes SMC foam cell formation by enhancing lipid uptake, we generated primary SMC explants from the *Adamts7*^SMCCTL^ and *Adamts7*^SMCTG^ mice with wild-type *Ldlr* and treated them with fluorescently labeled oxLDL (DiI-oxLDL). We chose oxLDL as it is a pathophysiologically relevant modified lipoprotein that accumulates within atherosclerotic lesions (22). *Adamts7*^SMCTG^ primary SMCs showed a 38% increase in DiI-oxLDL uptake compared to *Adamts7*^SMCCTL^ as measured by median fluorescence intensity (MFI) (Figure 4G). Both the proportion of DiI-oxLDL positive cells and the intensity of staining per cell were elevated in the *Adamts7*^SMCTG^ group. To further validate the role of *Adamts7* in SMC lipid uptake, we examined primary SMCs isolated from our previously described global *Adamts7* knockout mice (7). Following TNFα stimulation for 72 hours to induce *Adamts7* expression, the SMCs were loaded with DiI-oxLDL. Knockout of *Adamts7* reduced the percentage of cells accumulating lipids and the amount of lipid uptake (Figure 4H), complementing the transgenic phenotype.

### *Adamts7* leads to an increase in lipid uptake gene expression in SMCs

To identify how ADAMTS7 mediates an increase in SMC lipid uptake, we performed bulk RNA-seq on SMCs explanted from *Adamts7*^SMCCTL^ and *Adamts7*^SMCTG^ mouse aortas ten days after transgene induction (Figure 5A). These cells were from mice with wild-type *Ldlr* expression and fed a chow diet, ensuring that any changes in gene expression are due specifically to increased *Adamts7* expression. Differential gene expression analysis revealed that *Adamts7*^SMCTG^ cells had increased expression of lipid uptake genes typically associated with macrophages, including *Cd36, Fabp5*, and *Trem2,* as well as the macrophage-associated marker *Adgre1*. Importantly, this represents a transcriptional elevation; these cells are not bona fide macrophages. These findings were validated in vivo by qRT-PCR of whole aorta RNA from *Adamts7*^SMCTG^ mice (Figure 5B). While the increase in macrophage-like gene expression indicates SMC phenotypic modulation, we did not observe decreases in canonical SMC contractile markers, either by RNA-seq or by qRT-PCR (Supplemental Figure 5A), even with oxLDL treatment (Supplemental Figure 5B). Similarly, the expression of *Klf4*, a marker typically associated with modulated SMCs remained unchanged even with oxLDL treatment (13). Ingenuity Pathway Analysis (IPA) on the differentially expressed gene set identified pathways enriched within the *Adamts7*^SMCTG^ SMCs typically ascribed to macrophages, such as phagosome formation and immune signaling (Figure 5C). As one of the hallmark functions of macrophages during atherosclerosis is perpetuating inflammation, we asked whether our *Adamts7*^SMCTG^ SMCs were more inflammatory. Cytokine analysis of conditioned media revealed no differences in IL1β, IL6, or TNFɑ secretion (Supplemental Figure 5C). These findings indicate that *Adamts7*^SMCTG^ SMCs are not inflammatory and any atherosclerosis phenotype is likely independent of canonical inflammatory pathways. Collectively, the RNA-seq and qRT-PCR data indicate that *Adamts7*^SMCTG^ SMCs acquire a gene signature typically associated with macrophages, with an enrichment of phagocytic and lipid-uptake genes, consistent with prior reports that modulated SMCs can upregulate macrophage processes (12, 23).

**Figure 5.**
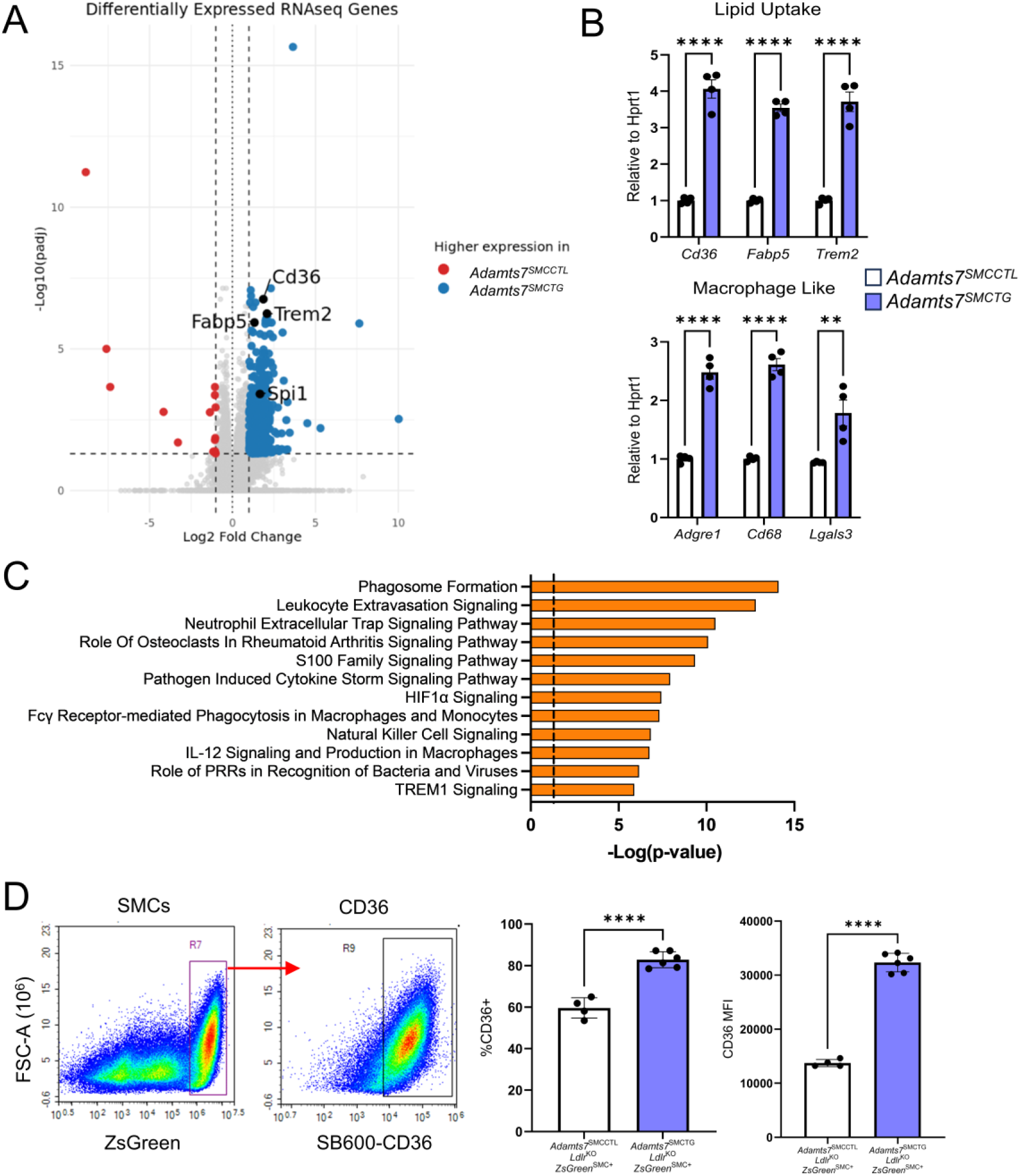
Bulk RNA-seq of *Adamts7*^SMCTG^ primary SMCs reveals an increase in lipid handling genes. (A) Volcano plot of the most upregulated and downregulated genes by *P* value adjusted for multiple comparisons. Passage two cells were used for RNA-seq. *Adamts7* (x = 3.65, y = 122) was excluded from the plot for visualization purposes because it fell completely off scale. (B) qRT-PCR confirmation of upregulated lipid uptake and macrophage-like genes *n =* 4. Normalization was performed relative to *Hprt1*. (C) Ingenuity Pathway Analysis of differentially expressed genes with Ingenuity Canonical Pathways highlighted. Shown pathways are predicted to be upregulated within *Adamts7*^SMCTG^ primary SMCs. (D) Confirmation of enhanced SMC CD36 in vivo. Flow cytometry analysis of lineage traced mice after 12 weeks of WTD feeding *n =* 4 - 6 mice. \*\*\*\**P*<0.0001, \*\**P*<0.01 Statistics were analyzed using a 2-tailed Student’s t-test.

### Knockdown of *Cd36* ameliorates enhanced oxLDL uptake conferred by *Adamts7*

Given the known role of CD36 as a receptor for oxLDL (24), we next investigated whether CD36 mediates the increased lipid uptake observed in *Adamts7*^SMCTG^ cells. Flow cytometric analysis was used on *Adamts7*^SMCTG^ *Ldlr*^KO^ ZsGreen^SMC+^ aortas after 12 weeks of WTD feeding to examine CD36 levels in vivo. ZsGreen^+^ cells from *Adamts7*^SMCTG^ *Ldlr*^KO^ ZsGreen^SMC+^ mice had a 40% increase in cells expressing *Cd36* and a 2.3-fold increase in MFI of CD36 signal (Figure 5D), consistent with the in vitro increase in CD36 expression in *Adamts7*^SMCTG^ cells. To determine whether CD36 is functionally required for ADAMTS7-induced lipid uptake, we performed siRNA-mediated knockdown of *Cd36* in primary SMCs isolated from *Adamts7*^SMCCTL^ and *Adamts7*^SMCTG^ mice. siRNA-mediated knockdown of *Cd36* in primary SMCs from *Adamts7*^SMCTG^ mice reduced *Cd36* expression to that of Non-Targeting Control (NTC) treated *Adamts7*^SMCCTL^ cells (Figure 6A). Consistent with this, siCd36 treatment normalized oxLDL uptake in *Adamts7*^SMCTG^ cells to levels comparable to NTC-treated controls (Figure 6B). Together, these data demonstrate that ADAMTS7 promotes SMC lipid uptake through upregulation of *Cd36*, identifying CD36 as a key mediator of the enhanced oxLDL uptake and foam cell formation driven by ADAMTS7 during atherogenesis.

**Figure 6.**
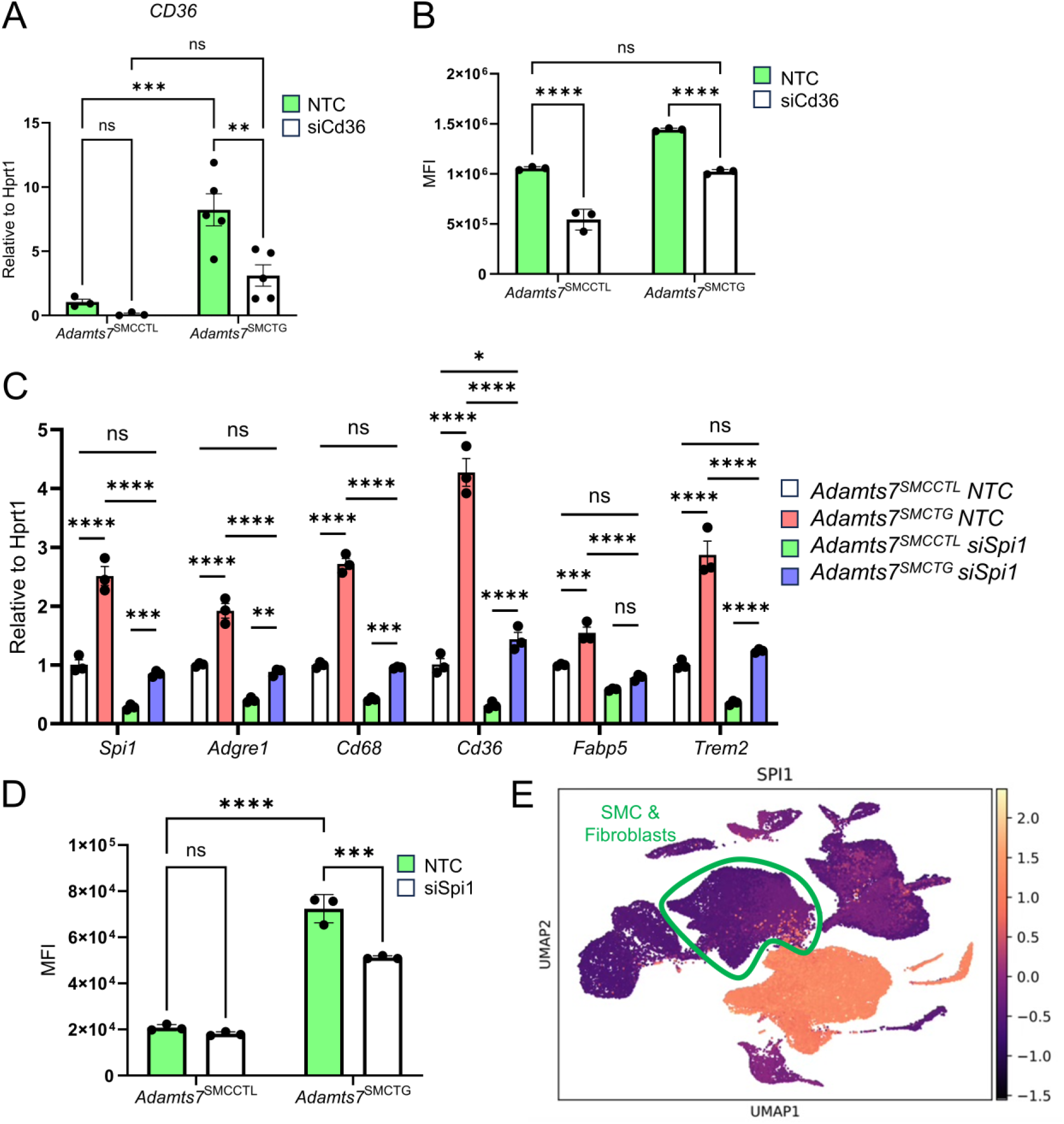
CD36 and PU.1 mediate ADAMTS7 conferred foam cell expansion. (A) Confirmation of knockdown of *Cd36* through siRNA. Primary SMCs were treated with 10 nM of either an NTC or *Cd36* siRNA. Knockdown was assessed 48 hours after transfection *n =* 3 - 5 mice. (B) Flow analysis of oxLDL lipid uptake. 24 hours post siRNA transfection, cells were treated with 10 µg/mL of DiI-oxLDL, and uptake was assessed 24 hours post oxLDL treatment *n =* 3. (C) Knockdown of *Spi1* in primary SMCs. Cells were treated with 10 nM of siRNA, and 48 hours after transfection, cells were harvested for qRT-PCR. (D) Flow analysis of lipid uptake. 24 hours post *Spi1* siRNA transfection, cells were treated with 10ug/mL of DiI-oxLDL, and uptake was assessed 24 hours post oxLDL treatment. (E) Expression of *SPI1* in human carotid atherosclerosis. \*\*\*\**P*<0.0001, \*\*\**P*<0.001, ** *P*<0.01, \**P*<0.05 Statistics were analyzed by two-way ANOVA with Sidak’s multiple comparison test.

### Knockdown of *Spi1* in *Adamts7* overexpressing SMCs attenuates the expression of macrophage-like and lipid-uptake genes

We next used IPA to identify upstream transcriptional regulators that could mediate the observed differential gene expression (Table 1) in *Adamts7*^SMCTG^ cells. The top predicted activated TF was *Spi1,* which encodes the protein PU.1. *Spi1* itself also exhibited a 3.16-fold increase in expression in *Adamts7*^SMCTG^ cells. To test whether PU.1 is required for the *Adamts7*-mediated increase in lipid uptake genes, we used siRNA to achieve a 66% knockdown of *Spi1* in primary SMCs. Knockdown of *Spi1* abrogated the increases in gene expression of *Cd36, Fabp5*, *Trem2, Adgre1*, and *Cd68* (Figure 6C). We also performed siRNA knockdown of other predicted master regulator TFs identified by IPA and found that knockdown of *Cebpa*, *Smarca4*, *Bhlhe40*, and *Klf6* did not ameliorate the increases in lipid uptake gene expression (Supplemental Figure 6). DiI-oxLDL loading of primary *Adamts7*^SMCTG^ SMCs with *Spi1* knockdown demonstrated that loss of *Spi1* attenuated the increased oxLDL uptake in *Adamts7*^SMCTG^ cells while having no effect in *Adamts7*^SMCCTL^ primary SMCs (Figure 6D). This finding suggests that increased *Spi1* levels resulting from *Adamts7* expression are responsible for the increased oxLDL uptake observed in *Adamts7*^SMCTG^ SMCs. Finally, we examined the level of *SPI1* expression within our human single-cell data and found clear *SPI1* expression in the modulated SMC populations (Figure 6E), mirroring our observations in mice and suggesting that *SPI1* may contribute similarly to SMC phenotypic modulation in humans. Within these human SMCs, phenotypic modulation was also accompanied by increased expression of lipid-handling genes *CD36* (19), *FABP5,* and *TREM2*, as well as the macrophage marker *ADGRE1* (Supplemental Figure 7).

**Table 1.** IPA upstream regulator analysis to identify candidate TFs. Upstream regulator analysis of RNA-seq reveals activated candidate molecules responsible for the gene expression changes. Five TFs are listed within the top 25 listed molecules.

| Upstream Regulator | Expr Fold Change | Molecule Type | Activation z-score |
| --- | --- | --- | --- |
| lipopolysaccharide | - | chemical drug | 5.491 |
| TNF | - | cytokine | 5.103 |
| E. coli B4 lipopolysaccharide | - | chemical toxicant | 4.872 |
| tetradecanoylphorbol acetate | - | chemical drug | 4.833 |
| **SPI1** | **3.16** | **transcription regulator** | **4.682** |
| CSF2 | - | cytokine | 4.675 |
| AGT | - | growth factor | 4.593 |
| OGA | - | enzyme | 4.593 |
| tretinoin | - | chemical drug | 4.484 |
| CNTF | - | cytokine | 4.243 |
| cardiotoxin | - | chemical - other | 4.123 |
| **BHLHE40** | - | **transcription regulator** | **4.022** |
| IL33 | - | cytokine | 3.943 |
| **SMARCA4** | - | **transcription regulator** | **3.939** |
| **CEBPA** | **3.969** | **transcription regulator** | **3.913** |
| D-glucose | - | chemical - endogenous mammalian | 3.907 |
| USP22 | - | peptidase | 3.898 |
| CD44 | - | other | 3.843 |
| poly rI:rC-RNA | - | biologic drug | 3.814 |
| IFNG | - | cytokine | 3.703 |
| ITPR2 | - | ion channel | 3.592 |
| TGM2 | - | enzyme | 3.568 |
| bleomycin | - | biologic drug | 3.567 |
| **KLF6** | - | **transcription regulator** | **3.549** |
| dihydrotestosterone | - | chemical - endogenous mammalian | 3.539 |

### ADAMTS7 promotes an AP-1-dependent chromatin remodeling program in SMCs

RNA-seq analysis revealed widespread transcriptional reprogramming in *Adamts7*^SMCTG^ SMCs (Figure 5), leading to increased oxLDL uptake. Given that ADAMTS7 is a secreted extracellular protein, these changes must be driven by outside-in signaling that alters TF activity. To identify changes in TF activity, we performed ATAC-seq on primary SMCs explanted from *Adamts7*^SMCCTL^ and *Adamts7*^SMCTG^ mice to map regions with altered chromatin accessibility. Differential ATAC-seq peak analysis revealed that *Adamts7* overexpression broadly remodels the chromatin landscape in SMCs, with 47,452 peaks (19.5% of all ATAC peaks) showing differential accessibility between groups (Figure 7A). TF binding motif analysis in the differential peaks identified enrichment of binding sites for FOS and JUN, members of the AP-1 TF complex (Figure 7B). Integration of ATAC-seq with RNA-seq data (Figure 5) further demonstrated that 85.8% of genes differentially expressed in *Adamts7*^SMCTG^ SMCs were associated with a nearby differential ATAC peak (±100 kb from the transcription start site), indicating a close coupling between chromatin accessibility and transcriptional reprogramming by ADAMTS7 (Figure 7C).

**Figure 7.**
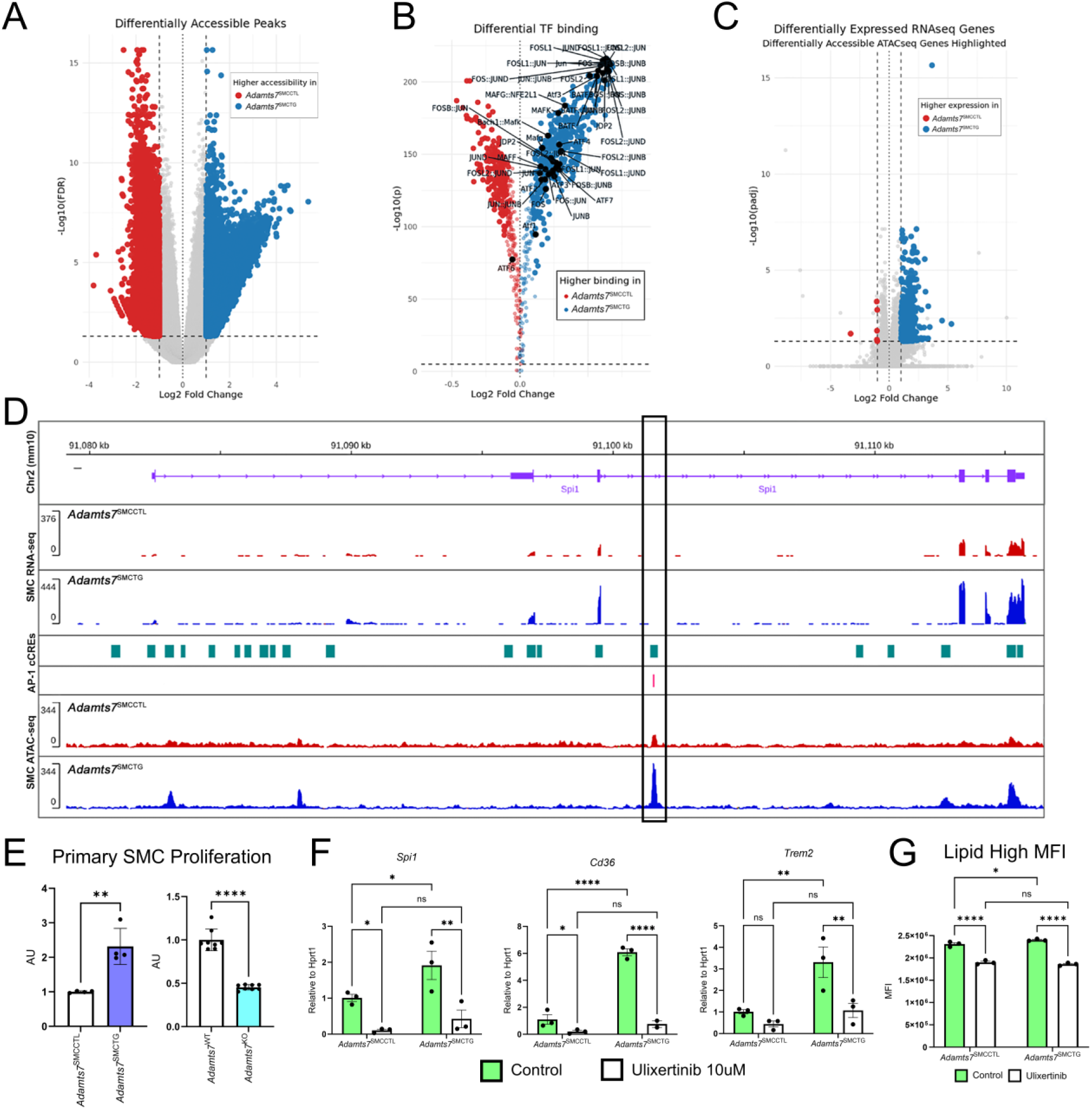
ADAMTS7 drives AP-1–dependent chromatin remodeling and transcriptional reprogramming in SMCs. (A) Differential ATAC-seq peaks in primary SMCs from *Adamts7*^SMCCTL^ and *Adamts7*^SMCTG^ mice reveal widespread chromatin remodeling. (B) Motif enrichment analysis of differential peaks identifies significant overrepresentation of FOS and JUN (AP-1) binding sites. (C) Integration of ATAC-seq and RNA-seq data shows that 85.8% of genes upregulated in *Adamts7*^SMCTG^ SMCs are associated with nearby differential ATAC peaks (±100 kb from TSS). (D) Genome browser tracks at the *Spi1* locus indicate increased chromatin accessibility in *Adamts7*^SMCTG^ SMCs at a predicted cis-regulatory element containing an AP-1 motif. (E) Proliferation as determined by CellTiter of primary SMCs of *Adamts7* overexpression *n =* 4 and *Adamts7* knockout cells *n =* 8. (F) qRT-PCR of ERK1/2 inhibition with Ulixertinib suppresses ADAMTS7-induced upregulation of *Spi1*, lipid-handling genes (*Cd36*, *Trem2*). (G) Ulixertinib treatment and subsequent DiI-oxLDL uptake in primary SMCs *n =* 3. Statistics were analyzed using a 2-tailed Student’s t-test or two-way ANOVA with Sidak’s multiple comparison test. \*\*\*\**P*<0.0001, \*\*\**P*<0.001, \*\**P*<0.01, \**P*<0.05

We next focused on the *Spi1* genomic locus in greater detail, chromatin accessibility increased markedly within ±200 kb of the *Spi1* transcriptional start site (Supplemental Figure 8). Notably, a region of differential accessibility within a *Spi1* intron overlaps both a predicted cis-regulatory element and AP-1 binding site, consistent with AP-1-dependent regulation of *Spi1* transcription in *Adamts7*^SMCTG^ SMCs (Figure 7D). AP-1 transcriptional activity is canonically regulated by MAPK/ERK-dependent phosphorylation of FOS and JUN, which enhances their DNA-binding and transactivation potential (25). Consistent with this model, *Adamts7*^SMCTG^ SMCs displayed increased proliferation, a hallmark of MAPK/ERK pathway activation, while SMCs isolated from global *Adamts7* knockout mice exhibited slower proliferation, confirming that ADAMTS7 contributes to SMC proliferative capacity (Figure 7E). We next tested whether inhibiting MAPK/ERK signaling could block ADAMTS7-induced phenotypes. Treatment with Ulixertinib, a selective ERK1/2 inhibitor, significantly suppressed *Spi1* expression (Figure 7F), reduced the induction of *Cd36* and *Trem2*, and blunted oxLDL lipid accumulation (Figure 7G). Together, these results are consistent with a model where ADAMTS7 promotes AP-1 activity, which in turn upregulates *Spi1* and downstream lipid-handling genes, driving both transcriptional and phenotypic reprogramming of SMCs.

Given that ADAMTS7 is a matrix metalloproteinase, we next investigated whether previously identified ADAMTS7 extracellular matrix (ECM) substrates (THBS1, COMP, LTBP4, and TIMP1) mediate the observed increases in lipid handling gene expression in *Adamts7*^SMCTG^ SMCs. To test this, we performed siRNA knockdowns of putative substrates in primary SMCs from *Adamts7*^SMCCTL^ and *Adamts7*^SMCTG^ mice (Supplemental Figure 9A). Among these, *Thbs1* knockdown most strongly attenuated the ADAMTS7-induced upregulation of *Cd36, Spi1*, and *Trem2*, while *Fabp5* expression remained unaffected. Although not fully restoring gene expression to baseline, this partial rescue indicates that *Thbs1* contributes in part to ADAMTS7-dependent gene activation. In contrast, *Comp* silencing modestly reduced the expression of select lipid-related genes with minor to no effects on *Cd36*, and knockdown of *Ltbp4* or *Timp1* produced minimal effects. Finally, we assessed THBS1 and COMP protein levels by western blot of whole aortic lysates following WTD feeding. This analysis revealed the presence of cleaved THBS1 fragments in *Adamts7*^SMCTG^ lysates with no reduction in total THBS1 protein, whereas total COMP protein levels were noticeably reduced (Supplemental Figure 9B). Collectively, these results demonstrate that ADAMTS7 promotes PU.1–dependent lipid uptake and transcriptional reprogramming in SMCs through a combination of ECM substrate–mediated signaling and AP-1 dependent chromatin remodeling.

## DISCUSSION

GWAS have revealed over 300 genomic regions associated with CAD (2, 26). Yet, functional studies elucidating the precise mechanisms by which these loci influence disease are still limited, hampering the clinical translation of these genetic findings. In 2011, our group and others reported *ADAMTS7* as a GWAS gene for CAD (4). Subsequent GWAS studies have replicated this signal multiple times and across different ethnic groups (2, 26), highlighting the importance of ADAMTS7 in CAD pathogenesis. Experimental studies in mice demonstrated that ADAMTS7 is proatherogenic and that its atherogenicity is conferred from its catalytic function (7, 9). Despite the sizable body of prior work on ADAMTS7, we still do not understand how ADAMTS7 participates in atherogenesis. Additionally, there have not been any cell-specific in vivo ADAMTS7 studies prior to this one, leaving the question of which vascular cell type(s) is critical for ADAMTS7 function. Here, we identified prominent expression of ADAMTS7 in human and mouse stromal and ECs and developed multiple novel mouse models in which *Adamts7* can be genetically overexpressed or conditionally knocked out in a cell-specific manner. Through these models, we demonstrate that both SMC- and EC-derived expression of *Adamts7* is proatherogenic, while knockout of *Adamts7* in either cell type alone is not sufficient to alter atherogenesis. These findings suggest that ADAMTS7 exerts redundant proatherogenic effects across multiple cell types through a shared mechanism in which it promotes SMC foam cell formation by enhancing oxLDL uptake via upregulation of *Spi1* and its downstream target *Cd36*.

*Adamts7* has very low overall expression across all murine vascular cell types in the basal state, and its expression is induced in response to vascular injury such as endothelial denudation or prolonged hyperlipidemia (7, 15). In primary mouse aortic SMCs, *Adamts7* induction is required to confer a detectable migration phenotype (7), confirming that this induction is critical for ADAMTS7 function. At terminal time points in mouse atherosclerosis studies, ADAMTS7 is again undetectable in plaques (7, 9), suggesting that *Adamts7* induction is transient and ADAMTS7 is mechanistically active in mice during plaque maturation. The transient induction of *Adamts7* has made studying its exact function in vascular cells difficult. The novel *Adamts7* mouse models presented here overcome this limitation, demonstrating that sustained *Adamts7* expression in either SMCs or ECs is sufficient to drive atherosclerosis progression.

While neither *Adamts7*^SMCKO^ *Ldlr*^KO^ nor *Adamts7*^ECKO^ *Ldlr*^KO^ affected atherosclerosis, prior research on whole-body knockout models indicates that *Adamts7* is pro-atherogenic. These prior published studies include three unique models of whole-body *Adamts7* inactivation. Two models employed genetic perturbation to inactivate ADAMTS7 by either exon deletion or mutation of the ADAMTS7 catalytic site (7, 9), while the third approach was to vaccinate against ADAMTS7 (10). While prior findings clearly demonstrate that *Adamts7* is expressed in cultured SMCs (7, 9, 15, 27), our RNAscope studies of *Adamts7* expression revealed its induction in multiple vascular cell types during murine atherosclerosis, yet only a small subset of vascular cells express *Adamts7* at any given time. Although we cannot exclude the existence of a yet unidentified dominant producer cell type, the reproducible multicellular distribution across independent datasets (19, 28) supports the view that *Adamts7* is broadly expressed and that its proatherogenic effects likely arise through cooperative or redundant mechanisms among vascular cells. Consistent with this, our data suggest that loss of *Adamts7* in a single cell type is insufficient to blunt atherogenesis, while broad inactivation across multiple compartments as in whole-body knockout reduces lesion burden (7, 9). Determining which combination of cell type–specific ADAMTS7 expression is required to drive atherosclerosis will necessitate future double-knockout models employing multiple Cre drivers.

Exactly how these findings translate to human atherosclerosis is another open question. Our analysis of the most extensive carotid plaque single-cell data set to date highlights that ECs and SMCs both express *ADAMTS7*, consistent with our murine findings and supporting the concept of multicellular expression in advanced disease. This scRNA-seq data also shows that mast cells and fibroblasts have detectable *ADAMTS7* expression, highlighting these cell types for further ADAMTS7 studies. Notably, the scRNA-seq data comes from mature atherosclerotic plaques, while our mouse data suggests the window for ADAMTS7 activity is earlier in plaque progression. Thus, whether this human expression pattern reflects relevant ADAMTS7 activity remains to be determined. Moreover, the therapeutic potential of targeting *ADAMTS7* in established lesions is unknown. Nonetheless, the human data indicate that ADAMTS7 is active in multiple vascular cell types, and our mouse data shows that sustained ADAMTS7 activity in individual cell types can drive atherogenesis. From a clinical perspective, much of the mechanistic understanding of ADAMTS7 function in atherosclerosis has been derived from murine models. Future studies examining functionally relevant *ADAMTS7* genetic variation, including the previously reported S214P common variant (27), as well as yet unidentified loss-of-function variants that impair catalytic activity, may help bridge mechanistic insights such as those described here with human disease pathology.

Our SMC-specific and EC-specific *Adamts7* overexpression mouse models have increased atherosclerotic burden and aortic foam cell formation, and these foam cells are SMC-derived. Although macrophages have historically been thought to be the source of foam cells in atherosclerotic lesions, it is increasingly appreciated that SMCs constitute most foam cells in atherosclerosis (14). Elegant fate mapping and integrative genomic studies have revealed that SMCs can transition into diverse phenotypic states, including inflammatory, fibroblast-like, and progenitor-like cells, through distinct transcriptional programs regulated by TFs such as KLF4, OCT4, TCF21, and retinoid signaling (11). Our findings raise the possibility that PU.1 is another TF directing SMCs specifically toward the foam cell fate. Consistent with this, when *Adamts7* is induced in the vasculature, SMCs increase their expression of genes involved in lipid uptake, inflammation, and phagocytosis, a gene expression profile reminiscent of a macrophage. As recent studies have definitively shown that SMCs do not contribute to the pool of mature macrophages in atherosclerosis (29), we posit that these *Adamts7*^SMCTG^ SMCs are modulated SMCs with increased lipid uptake capacity and foam cell features. While the specific role of SMC foam cells in atherosclerosis remains unclear, it has been reported that SMC foam cells and true macrophages differ in their intracellular lipid metabolism (30, 31). Lipid-laden, foamy macrophages tend to be less inflammatory (32), and by becoming foamy, these macrophages may no longer contribute to the inflammatory cycle associated with atherosclerosis, raising the possibility that foamy macrophages are beneficial in atherosclerosis. Conversely, we observed reduced fibrous cap thickness with *Adamts7* overexpression. As one well-described role of SMCs is forming the fibrous cap, it is possible that foamy SMCs cannot stabilize lesions, raising an intriguing possibility that macrophage foam cells are beneficial, whereas SMC foam cells are detrimental. In Sharifi et al., the authors showed that ADAMTS7 degrades TIMP1, increasing MMP2 and MMP9 activity (33). Consistent with that model, ADAMTS7 whole-body knockout increased collagen content and plaque stabilization, and *ADAMTS7* expression was found to be higher in caps of unstable human carotid plaques (33). These prior observations and those presented here support the adverse role of ADAMTS7 in plaque stability and suggest multiple mechanistic explanations for ADAMTS7-mediated plaque instability.

Mechanistically, we chose to pursue the function of SMC *Adamts7* given the large body of preexisting data on *Adamts7* modulation of SMC function. We find that constitutive *Adamts7* SMC expression markedly alters lipid uptake gene programs, resulting in enhanced oxLDL accumulation. Among these lipid uptake genes, one of the most significantly upregulated genes was *Cd36*. CD36 is a well-described receptor for oxLDL, and our studies demonstrate that CD36 is mainly responsible for the SMC foam cell phenotype in *Adamts7*^SMCTG^ mice (24). In addition, upstream regulator analysis of primary SMCs identified *Spi1* as a potential mediator of expression changes (34), and knockdown of *Spi1* in *Adamts7*^SMCTG^ SMCs ameliorated the increase in lipid-handling gene expression and macrophage markers. *Spi1*, which encodes the TF PU.1, is well recognized as a critical mediator of hematopoietic lineage commitment. These observations suggest that PU.1 may also be a crucial factor in driving the in vivo production of foamy SMCs. Thus, future cell-specific studies of SMC PU.1 may help reveal disease-relevant consequences of foamy SMCs.

Using ATAC-seq, we observed that ADAMTS7 enhances chromatin accessibility and activates AP-1–dependent transcription. AP-1 activity is regulated by MAPK signaling via ERK1/2, and pharmacologic inhibition of ERK1/2 reduced ADAMTS7-induced expression of lipid-handling genes. These findings support a model in which extracellular ADAMTS7 activates MAPK-ERK1/2 signaling, modulating AP-1 activity and altering SMC gene expression profiles. Importantly, this suggests that ADAMTS7-mediated cleavage of an extracellular substrate triggers MAPK signaling cascades. ADAMTS7 substrates have been proposed in prior studies (8, 33, 35, 36), but none have been tested in the context of SMC foam cell formation. To explore this, we examined several previously proposed substrates and found that *Thbs1* knockdown partially attenuated ADAMTS7-associated lipid gene expression. Western blot analysis of aortic lysates revealed cleaved THBS1 fragments without a reduction in total THBS1 protein, while COMP levels were modestly reduced, suggesting differential substrate processing in vivo. Notably, *Comp* silencing also suppressed *Spi1* expression, raising the possibility that COMP may contribute to ADAMTS7-mediated oxLDL uptake. Together, these findings support a model in which ADAMTS7-dependent substrate processing, potentially involving both THBS1 and COMP, contributes to downstream signaling. The identification of ADAMTS7 cleavage substrates remains an active area of investigation. Although THBS1 and COMP have been previously reported as ADAMTS7 substrates (17, 37), not all studies have reproduced these findings (35), and the relative contribution of these substrates remains undefined. Definitive evaluation of these substrates is necessary, as identifying biologically meaningful cleavage products may reveal functional therapeutic biomarkers and inform the development of targeted ADAMTS7-directed therapies.

In conclusion, our study establishes that ADAMTS7 promotes SMC foam cell formation and atherosclerosis through PU.1 and CD36 and that its expression is not confined to a single vascular cell type. By integrating human single-cell data with new cell-specific genetic models, we show that ADAMTS7 is distributed across diverse vascular populations and that its sustained activation, whether from SMCs or ECs, drives disease progression. These findings reveal a unifying mechanism linking human GWAS data with cellular pathophysiology, reinforcing ADAMTS7 inhibition as a potential therapeutic target for atherosclerotic cardiovascular disease.

## METHODS

### Sex as a biological variable

Our human data contained both male and female samples. SMC mouse studies were limited to male mice due to the integration of the *Myh11*-CreER^T2^ onto the Y-chromosome.

### Animals

C57BL/6J (Stock no. 000664), *Ldlr*^KO^ (Stock no. 002207), ZsGreen (Stock no. 007906), and *Myh11*-CreER^T2^ (Stock no. 019079) were purchased from the Jackson Laboratory and bred in our laboratory. *Adamts7* knockout mice were previously described (7). The *Cdh5*-CreER^T2^ Mouse was a gift from Dr. Carol Troy (Columbia University, New York City, USA) and developed by the laboratory of Dr. Ralf Adams (Max Planck Institute for Molecular Biomedicine, Münster, Germany) and has been previously described (38). To generate a floxed *Adamts7*, the *Adamts7* knockout mouse (MMRRC, 046487) was bred to an FLP recombinase mouse. *Adamts7* transgenic mice were created by cloning the mouse allele of *Adamts7* into the pR26 CAG/GFP Asc plasmid (Addgene, plasmid # 74285). Subsequently, FL19 embryonic stem cells were used to generate chimeras, which were then backcrossed to the C57BL/6J background. For CreER^T2^ activation, tamoxifen (Sigma-Aldrich, T5648) was dissolved in 90% corn oil and 10% ethanol; subsequently, mice at seven weeks of age were injected with tamoxifen for five days at a dosage of 40 mg/kg of body weight per day. At eight weeks of age, the mice were fed a WTD consisting of 40% Kcal by fat and 0.15% Cholesterol (Research Diet, D12079Bi). For blood collection, mice were fasted for four hours and subsequently sedated with isoflurane, and blood was collected by retroorbital puncture with heparinized microcapillary tubes. Plasma was isolated by centrifugation at 1500 RCF for 10 minutes at 4°C. Total cholesterol was measured using the Infinity Cholesterol Reagent (Thermo Fisher Scientific, TR13421).

### Atherosclerosis analysis

For atherosclerosis studies, mice were sacrificed by cervical dislocation and perfused with PBS. The aorta was dissected and subsequently fixed in 10% formalin. Whole aortas were stained with ORO. Hearts containing the aortic root were fixed in 4% paraformaldehyde for 24 hours, transferred to 70% ethanol, and embedded in paraffin wax. Sectioning was performed using a Leica microtome at 8 µm thickness. The heart was sectioned from the inferior heart towards the aortic root. 25 slides were collected, with each slide containing two sections. We aimed to have the breakage of the leaflet occur on slide 13. Slides 3, 8,13,18, and 23 were used for hematoxylin and eosin staining to allow for lesion size quantification using ImageJ. Hematoxylin and Eosin, and Picrosirius Red staining was performed at Columbia’s Molecular Pathology Shared Resource.

### Immunofluorescence

Immunofluorescence was performed on paraffin sections of the aortic root. Deparaffinization was performed using xylene. Antigen retrieval was performed using a citrate-based agent. Sections were subsequently blocked using 10% normal goat serum for at least one hour. Subsequently, the sections were stained using the following antibodies and dilutions 1:1000 αSMA-Cy3 (Sigma Aldrich, C6198), 1:500 MAC2 (Cedarlane, CL8942AP). IgG controls were purchased from Jackson Immunoresearch. The sections were washed with PBS containing 0.01% Tween 20. For the secondary antibody, a donkey anti-rabbit Alexa Fluor 647 (Invitrogen, A-31573) and goat anti-mouse Alexa Fluor 546 (Invitrogen, A-11030) were used, both at a 1:1000 dilution in 1% goat serum. The slides were washed with DAPI and mounted with ProLong Glass Antifade Mountant with NucBlue Stain. Imaging was conducted on a Nikon Eclipse Ti-S microscope, and image processing was performed using ImageJ. ZsGreen fluorescence was confirmed separately on unfixed frozen sections without additional antibody staining to verify reporter expression.

### Vascular SMC isolation for tissue culture

SMCs were isolated as previously described (39). The aorta was dissected from the ascending arch to the diaphragm and was predigested for 10 minutes at 37°C with 175 U/mL Collagenase II and 1.25 U/mL elastase in HBSS to allow for the removal of the adventitia. After removing the adventitia, the media layer was further digested in 400 U/mL collagenase II, 2.5 U/mL elastase, and 0.2 mg/mL soybean trypsin inhibitor at 37°C for one hour in a rotating incubator. The cells were cultured in DMEM supplemented with 20% fetal bovine serum (FBS), 1% L-Glutamine, 1% sodium pyruvate, and 1% Penicillin Streptomycin. Cells between passages two and seven were used for experiments.

### Vascular SMC isolation for flow analysis

Mice were sacrificed and perfused with PBS. The ascending aorta to the bifurcation at the common iliac artery was dissected and digested with a cocktail consisting of 4 U/mL Liberase TM (Sigma-Aldrich, 5401127001), 60 U/mL hyaluronidase (Sigma-Aldrich, H3506), and 60 U/mL DNase I (Worthington Biochemical Corporation, LS006333) in RPMI-1640. The digestion enzymes were neutralized with 10% FBS in RPMI and washed once in FACS buffer (2% FBS, 5 mM EDTA, 20 mM HEPES, and 1 mM Sodium Pyruvate in DPBS). Cells were then blocked with TruStain FcX PLUS anti-mouse CD16/32 (Biolegend, 156603) for 10 mins at 4°C. Staining was performed in FACS buffer at the following dilutions: 1:1000 LipidTOX Red (Thermo Fisher Scientific, H34476), 1:100 CD36 Super Bright 600 (Invitrogen, 63-0362-82), 1:200 CD45-Alexa Fluor 700 (Biolegend, 157209). Cell viability was assessed with DAPI. To establish the LipidTOX high population and thus foamy cells, a C57BL/6J mouse with normal lipidemia was used as previously described (14).

### In vitro cell culture experiments

For in vitro foam cell analysis, cells were treated with DiI conjugated oxLDL (Invitrogen, L34358) at a concentration of 10 µg/mL for 24 hours. Subsequently, they were washed with PBS and detached using 0.25% Trypsin EDTA. Neutralization of trypsin was performed using FACS buffer. The cells were counterstained with DAPI to assess live cells, and all analysis was done using either a Novocyte Quanteon or Penteon. For siRNA experiments, all siRNAs were purchased from Integrated DNA Technologies (Supplemental Table 2). Transfection was performed using RNAiMAX according to the manufacturer’s protocol. All siRNAs were used at a concentration of 10 nM.

### RNA Isolation and quantitative RT-PCR

For tissue samples, specimens were collected in Trizol, and homogenization was performed using a tissue lyser (Qiagen). The RNA was extracted using the Direct-zol RNA MiniPrep Kit (Zymo) according to the manufacturer’s instructions. For cells, RNA was collected using the Quick-RNA Microprep Kit (Zymo). Complementary DNA was synthesized using High-Capacity cDNA Reverse Transcription Kit (Applied Biosystem). All qPCR was performed using TaqMan probes (Supplemental Table 2) and performed on the Applied Biosystems QuantStudio 7 Flex Real-Time PCR System. Analysis was performed using the 2-ΔΔCt Method.

### RNAscope

RNAscope was performed according to the manufacturer’s instructions. RNAscope was performed on sections of mouse BCAs after specified durations of WTD feeding. The BCAs were freshly frozen and sectioned at 8 µm thickness. The sections were first fixed in 10% neutral buffered formalin. Subsequently, the sections were dehydrated and treated with hydrogen peroxide and protease. Hybridization with *Adamts7* (Advanced Cell Diagnostics, 533341) and *Pecam1* (Advanced Cell Diagnostics, 316721-C3) probes was performed. After probe hybridization, hybridization with AMPs 1-3 was performed. Signal development was conducted using Akoya Biosciences Opal Reagent Kits, and slides were mounted using Prolong Glass Antifade Mountant (Thermo Fisher Scientific, P36984). Imaging was conducted on a Nikon Eclipse Ti-S microscope, and image processing was performed using ImageJ.

### Western blot

Samples were lysed in RIPA buffer (Thermo Fisher Scientific, 89900) supplemented with 1% Halt Protease and Phosphatase Inhibitor Cocktail. Tissue samples were lysed with the Qiagen TissueLyser and samples were subsequently cleared by centrifugation at 20,000 RCF for 10 minutes at 4°C. Samples were denatured, resolved on a 4-12% Bis-Tris gel, and transferred to either PVDF or a nitrocellulose membrane. Blocking was performed with 5% milk, and primary antibodies were diluted 1:1000 in 5% milk and incubated overnight at 4°C. The following primary antibodies were used: β-Actin (Cell Signaling Technology, 5125S), Thrombospondin-1 (D7E5F) Rabbit mAb (Cell Signaling Technology, 37879S), GAPDH (14C10) Rabbit mAb, HRP-conjugated (Cell Signaling Technology, 3683S), Anti-COMP/Cartilage Oligomeric Matrix Protein (Abcam, ab42225), and Rabbit Ab ADAMTS7 (custom-made by 21st Century Biochemicals). The custom polyclonal rabbit anti–ADAMTS7 antibody was generated using standard immunization protocols with recombinant mouse ADAMTS7 as the antigen. Membranes were washed with 0.1% TBST and probed with the appropriate secondary antibody for one hour at room temperature. Membranes were visualized with ECL using the Amersham Imager 600 gel imager.

### RNA-seq of primary mouse vascular SMCs

Primary SMCs were isolated as described above and cultured for three passages spaced one week apart. Total RNA was extracted using the Zymo Microprep Kit, and RNA integrity was assessed using TapeStation. No cell sorting was performed. Polyadenylated mRNA was purified from total RNA using a poly(A) pull-down and reverse-transcribed into cDNA, with the final PCR step performed using KAPA HiFi HotStart Ready Mix. Libraries were sequenced on the Element AVITI sequencer with 75 bp paired-end reads, targeting ∼40 million reads. Transcript abundance was quantified using kallisto with the GRCm38 mouse transcriptome, and differential expression analysis was performed using DESeq2 via DEBrowser. Up and downregulated genes are listed in Supplemental Table 1.

### ATAC-seq

Assay for Transposase-Accessible Chromatin using sequencing (ATAC-seq) was performed as previously described with minor modifications. Briefly, 100,000 passage 3 primary mouse SMCs with >90% viability were washed with PBS and pelleted at 500 × g for 5 min at 4°C. Cells were lysed in cold lysis buffer (10 mM Tris-HCl pH 7.4, 10 mM NaCl, 3 mM MgCl₂, 0.1% NP-40, 0.1% Tween-20, and 0.01% digitonin). Nuclei were washed with 1 mL ATAC lysis buffer without NP-40 or digitonin and centrifuged at 500 × g for 10 min at 4°C. The nuclei pellet was resuspended in transposition mix containing 25 μL 2× TD buffer (Illumina Nextera DNA Library Prep Kit), 2.5 μL Tn5 transposase, 16.5 μL PBS, 0.5 μL 1% digitonin, 0.5 μL 10% Tween-20, and 5 μL nuclease-free water. Transposition was carried out at 37°C for 30 min in a thermomixer. DNA was purified using the Zymo DNA Clean & Concentrator-5 Kit. Transposed DNA was amplified using NEBNext High-Fidelity 2× PCR Master Mix (New England Biolabs, M0541) with Nextera indexing primers, and libraries were purified with AMPure XP magnetic beads (Beckman Coulter, A63880). Library size distribution and quality were assessed using an Agilent Bioanalyzer, and DNA concentration was measured with a Qubit dsDNA HS Assay Kit (Thermo Fisher Scientific). Paired-end 150 bp sequencing was performed on an Illumina platform by Azenta/Genewiz. Sequencing reads were quality-trimmed with Trimmomatic v0.38, aligned to the mm10 reference genome using Bowtie2, and filtered for uniquely mapped, high-quality (MAPQ ≥30) non-duplicate reads with SAMtools v1.9 and Picard v2.18.26. Reads mapping to mitochondrial DNA and unplaced contigs were excluded. Peak calling was performed using MACS2 v2.1.2, with peaks overlapping ENCODE blacklisted regions removed. Peaks detected in ≥66% of biological replicates per group were retained for downstream analysis. Differential chromatin accessibility was determined using DiffBind (R package). Sequencing yielded a total of 528 million paired-end reads (mean Q score = 36.9; 85% bases ≥ Q30). After alignment (mean alignment rate = 98%) and filtering, approximately 60–100 million high-quality read pairs per sample were retained for downstream analyses.

### Statistics

All statistical tests are indicated in figure legends. *P* values were calculated using a 2-tailed Student’s t-test in GraphPad Prism 10. *P*<0.05 was considered significant. 2-way ANOVA analysis with multiple corrections was used for comparison between groups with post-analysis as indicated in the figure legends. Aortic root histology, qPCR, and ELISA results are presented as mean ± standard error of the mean. All other data are presented as mean ± standard deviation.

### Study approval

All mouse experiments were performed according to procedures approved by Columbia University’s Institute for Animal Care and Use Committee under protocol AABU8650. The Institutional Review Board of Columbia University approved all human studies under IRB AAAJ2765 and AAAR6796 with written informed consent provided by all participants.

## Data availability

Human carotid artery scRNA-seq has been previously published (19). ATAC and RNA-seq data have been deposited in the NCBI’s Gene Expression Omnibus database (GSE316135). All individual values represented in graphs are provided in the Supporting Data Values file.

## AUTHOR CONTRIBUTIONS

A.C. led the study and directly performed the majority of experiments. H.K.C., K.S., C.V.M., and J.G. P. conducted experiments and acquired data. H.P. contributed to the generation of mouse lines. L.E.F. and J.S.K. performed ATAC-seq data analysis. A.C.B., H.Y., and M.L. provided human single-cell datasets. R.C.B. oversaw study design, execution, management, and funding. A.C. and R.C.B. wrote the manuscript, with all authors reviewing and approving the final version.

## ACKNOWLEDGMENTS

We thank Dr. Muredach P. Reilly for guidance in the execution of these studies and Dr. Elizabeth E. Ha for helpful discussions. We acknowledge the staff of the Columbia Stem Cell Initiative Flow Cytometry Core Facility, led by Michael Kissner at Columbia University Irving Medical Center, for their technical support. We also thank Dr. Carol Troy for generously providing the *Cdh5*-CreER^T2^ mice.

