## Supplemental Data for "ADAMTS7 promotes smooth muscle foam cell expansion in atherosclerosis"

### SUPPLEMENTAL FIGURE 1

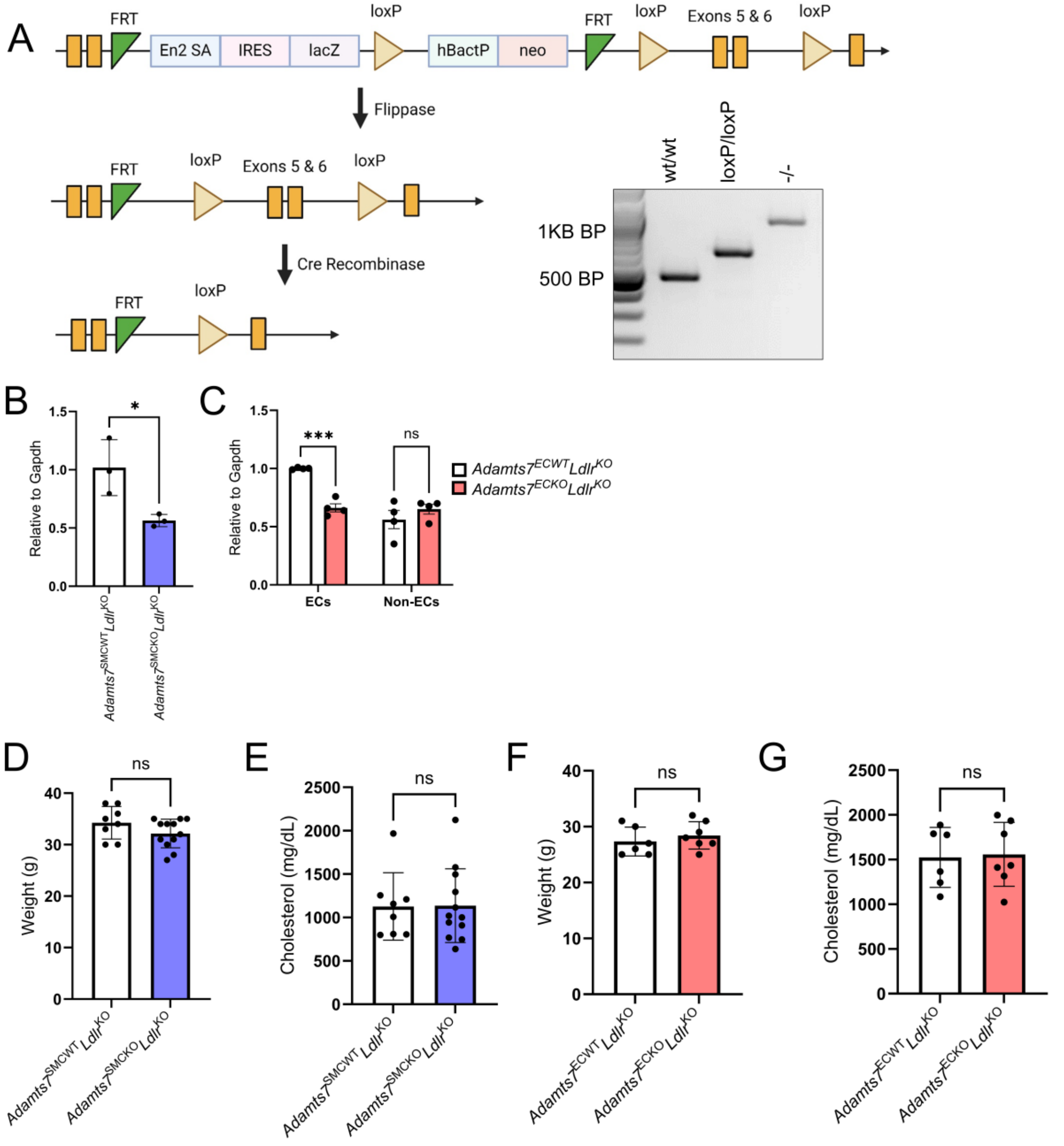

**Supplemental Figure 1. Design of conditional knockout mice and weights and cholesterol after WTD.** (A) Schematic overview of the knockout-first allele. Knockout mice were crossed with the flp recombinase deleter mice, generating mice with floxed *Adamts7* exons 5 and 6. Subsequently, breeding the conditional knockout mice with a Cre recombinase produces tissue-specific *Adamts7* knockout. Genotyping of wild-type, conditional knockout, and whole-body knockout mice. (B) Primary SMCs were explanted from *Adamts7*$^{SMCWT}$ *Ldlr*$^{KO}$ and *Adamts7*$^{SMCKO}$ *Ldlr*$^{KO}$ mice following tamoxifen induction and analyzed for *Adamts7* expression ($n$ = 3 mice). (C) Verification of *Adamts7* knockout in CD31 bead-purified mouse aortic endothelial ($CD31^+$) and non-endothelial ($CD31^-$) cells ($n$ = 4). (D) Mice's weight (grams) and (E) plasma total cholesterol after 16 weeks of WTD feeding for *Adamts7*$^{SMCWT}$ *Ldlr*$^{KO}$ and *Adamts7*$^{SMCKO}$ *Ldlr*$^{KO}$ mice. $n$ = 8 - 12 male mice. (F) Mice's weight (grams) and (G) plasma total cholesterol after 16 weeks of WTD feeding for *Adamts7*$^{ECWT}$ *Ldlr*$^{KO}$ and *Adamts7*$^{ECKO}$ *Ldlr*$^{KO}$ mice. $n$ = 6 - 7 female mice. Statistics were analyzed using a 2-tailed Student's t-test. *$P < 0.05$; ***$P < 0.001$

SUPPLEMENTAL FIGURE 2

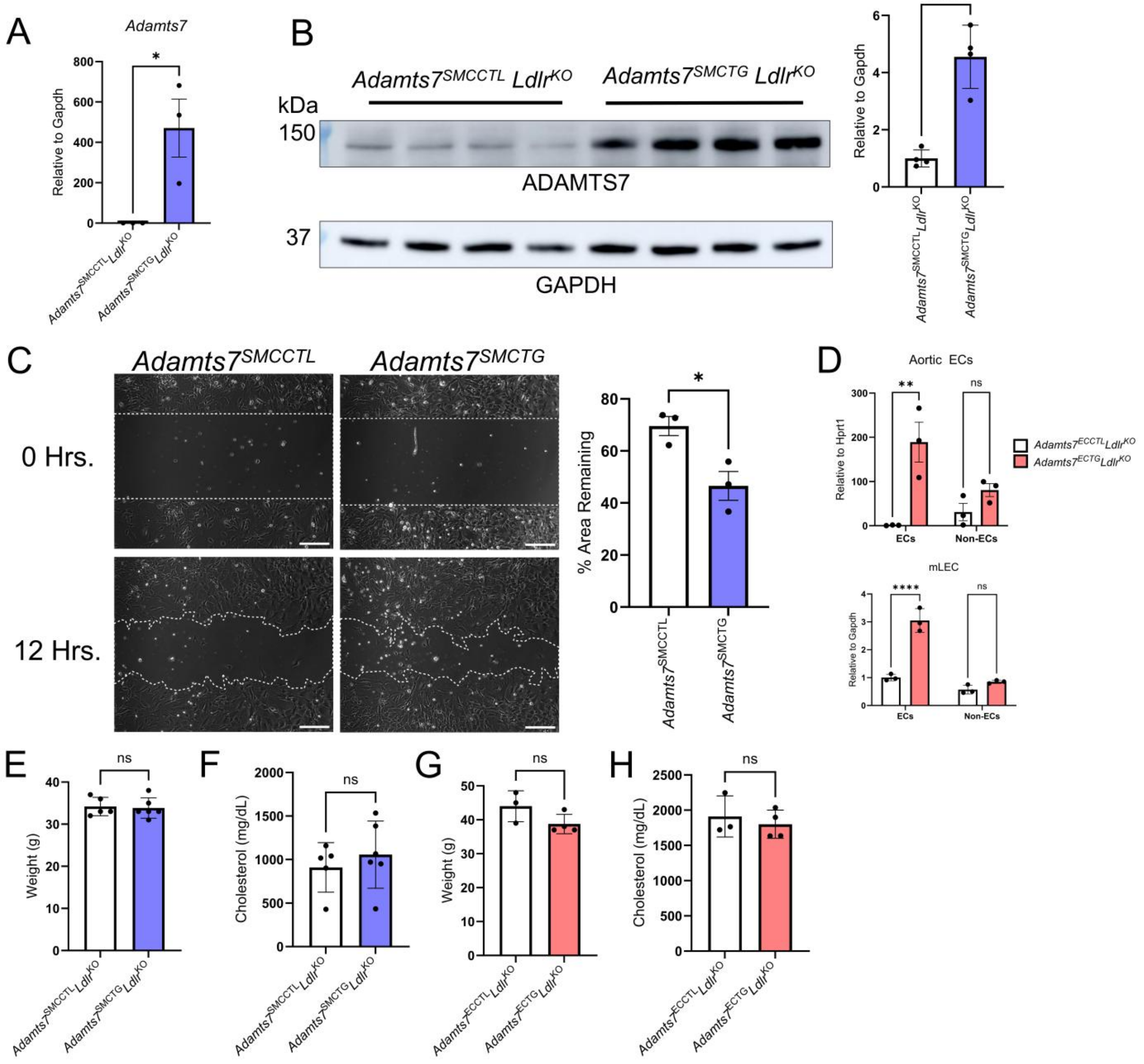

**Supplemental Figure 2. Verification and validation of transgenic mice**. (A) Verification of *Adamts7* overexpression in $Adamts7^{SMCTG}$ $Ldlr^{KO}$ mice by qPCR *n* = 3 male mice. (B) Validation of ADAMTS7 protein overexpression *n* = 4 male mice. (C) $Adamts7^{SMCTG}$ mice have enhanced migration. *n* = 3 male mice. Scale bar = 200 μm (D) Confirmation of *Adamts7* overexpression in bead-purified aortic endothelial ($CD31^{+}$) and non-endothelial ($CD31^{-}$) cells from EC-Adamts7 transgenic mice (*n* = 3 mice). Verification of *Adamts7* transgenic expression in isolated mouse lung ECs ($CD31^{+}$) and non-ECs ($CD31^{-}$) *n* = 3 mice. (E) Mice's weight (grams) and (F) plasma total cholesterol after 16 weeks of WTD feeding for $Adamts7^{SMCCTL}$ $Ldlr^{KO}$ and $Adamts7^{SMCTG}$ $Ldlr^{KO}$ mice. *n* = 5 - 7 male mice. (G) Mice's weight (grams) and (H) plasma total cholesterol (mg/dL) after 16 weeks of WTD feeding for $Adamts7^{ECCTL}$ $Ldlr^{KO}$ and $Adamts7^{ECTG}$ $Ldlr^{KO}$ mice. *n* = 3 - 4 female mice. Statistics were analyzed using a 2-tailed Student's t-test. ****$P$<0.0001, ***$P$<0.001, **$P$<0.01, *$P$<0.05

### SUPPLEMENTAL FIGURE 3

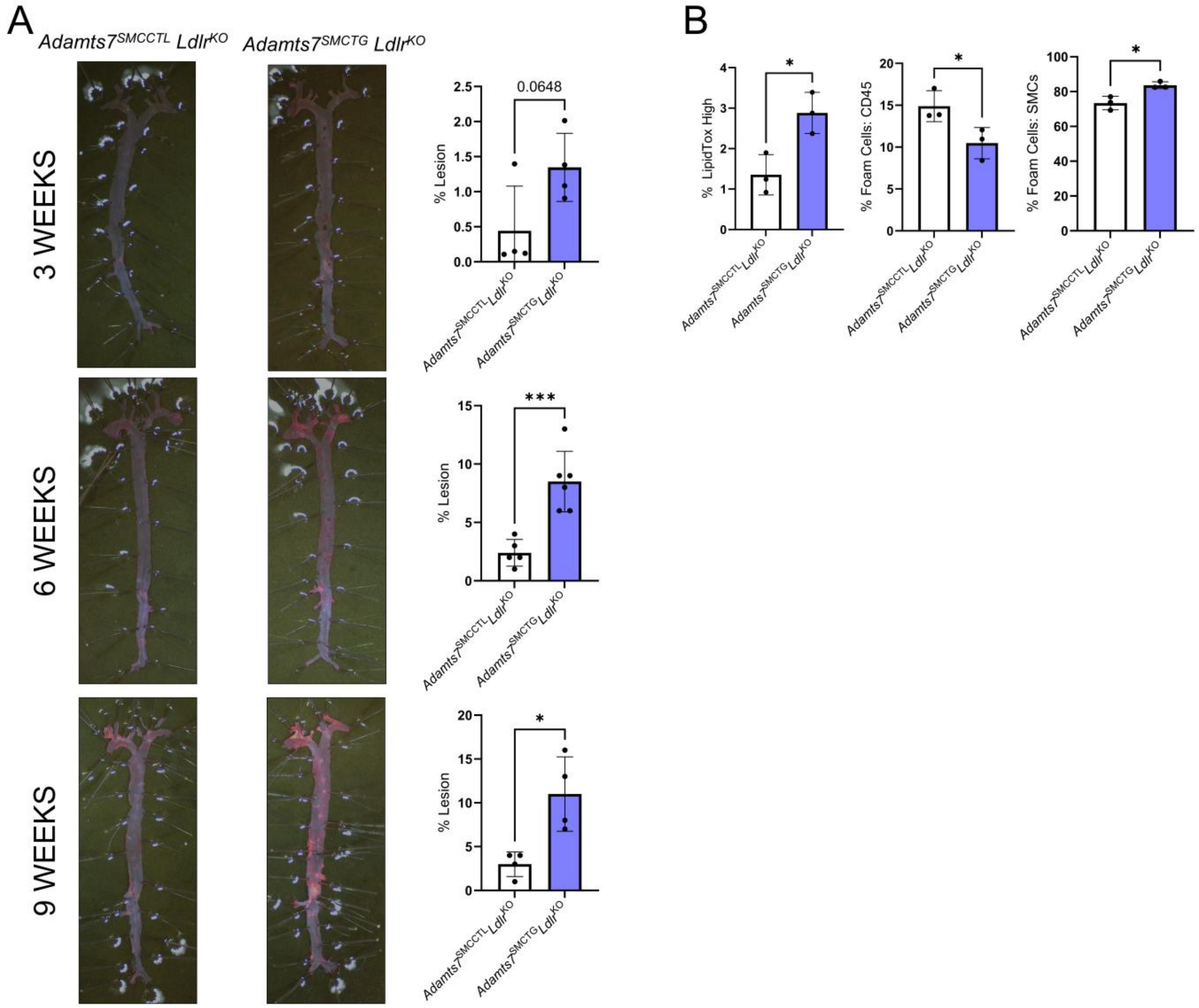


**Supplemental Figure 3. Increased atherosclerosis with short durations of WTD feeding in *Adamts7*$^{\mathbf{SMCTG}}$ mice.**

(A) Representative images of ORO stained en-face aortas after three, six, and nine weeks of WTD feeding. *n* = 4 - 6 male mice (B) Foam cell analysis of aortas after 3 weeks of WTD feeding. Aortic single-cell suspensions were stained for leukocyte marker CD45 and smooth muscle cell marker CD200 *n* = 3 male mice. *$P<0.05$; ***$P<0.001$ Statistics were analyzed using a 2-tailed Student's t-test.

SUPPLEMENTAL FIGURE 4

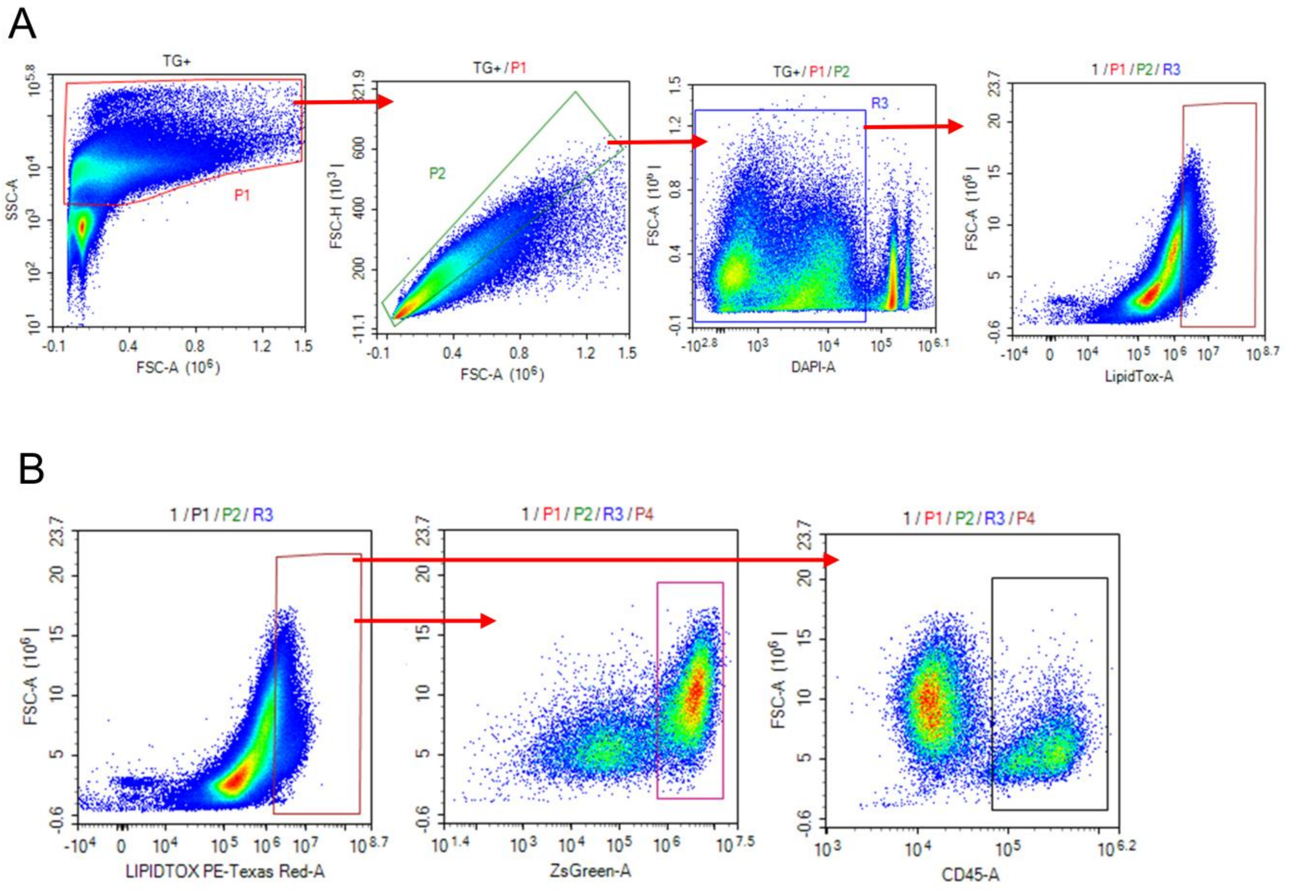


**Supplemental Figure 4. Representative gating strategy for foam cells.**

(A) Representative flow cytometry gating strategy. Cells were isolated from debris. Singlets and live cells (DAPI negative) were subsequently analyzed. LipidTOX high gate was established using a normal lipidemic C57BL/6J aorta. (B) LipidTOX high foam cells were further assessed for ZsGreen expression and stained for the leukocyte marker CD45.

#### SUPPLEMENTAL FIGURE 5

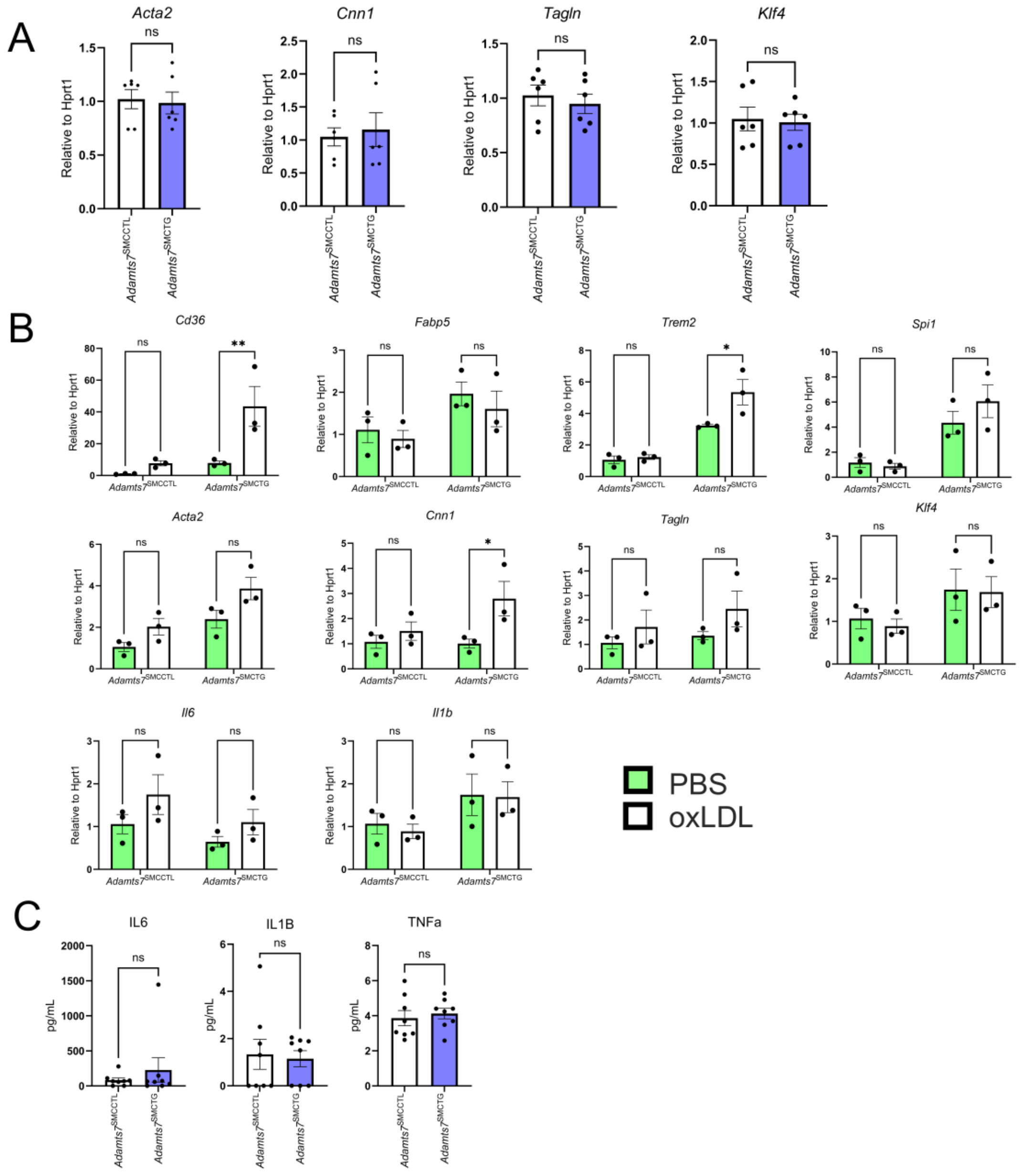


**Supplemental Figure 5. Characterization of explanted primary SMCs**

(A) qPCR characterization of contractile and modulated genes. *n* = 6. (B) SMCs explanted from *Adamts7*$^{\mathrm{SMCCTL}}$ and *Adamts7*$^{\mathrm{SMCTG}}$ mice were treated with 100 µg/mL oxLDL for 72 hours, followed by qRT-PCR analysis of the indicated genes. (C) LEGENDplex-based ELISA is used to detect inflammatory cytokines in cell culture-conditioned media. Cells were from *n* = 8 mice. Statistics were analyzed using a 2-tailed Student's t-test or two-way ANOVA with Sidak's post hoc test for multiple comparisons. *****P*<0.0001, ****P*<0.001, ***P*<0.01, **P*<0.05

### SUPPLEMENTAL FIGURE 6

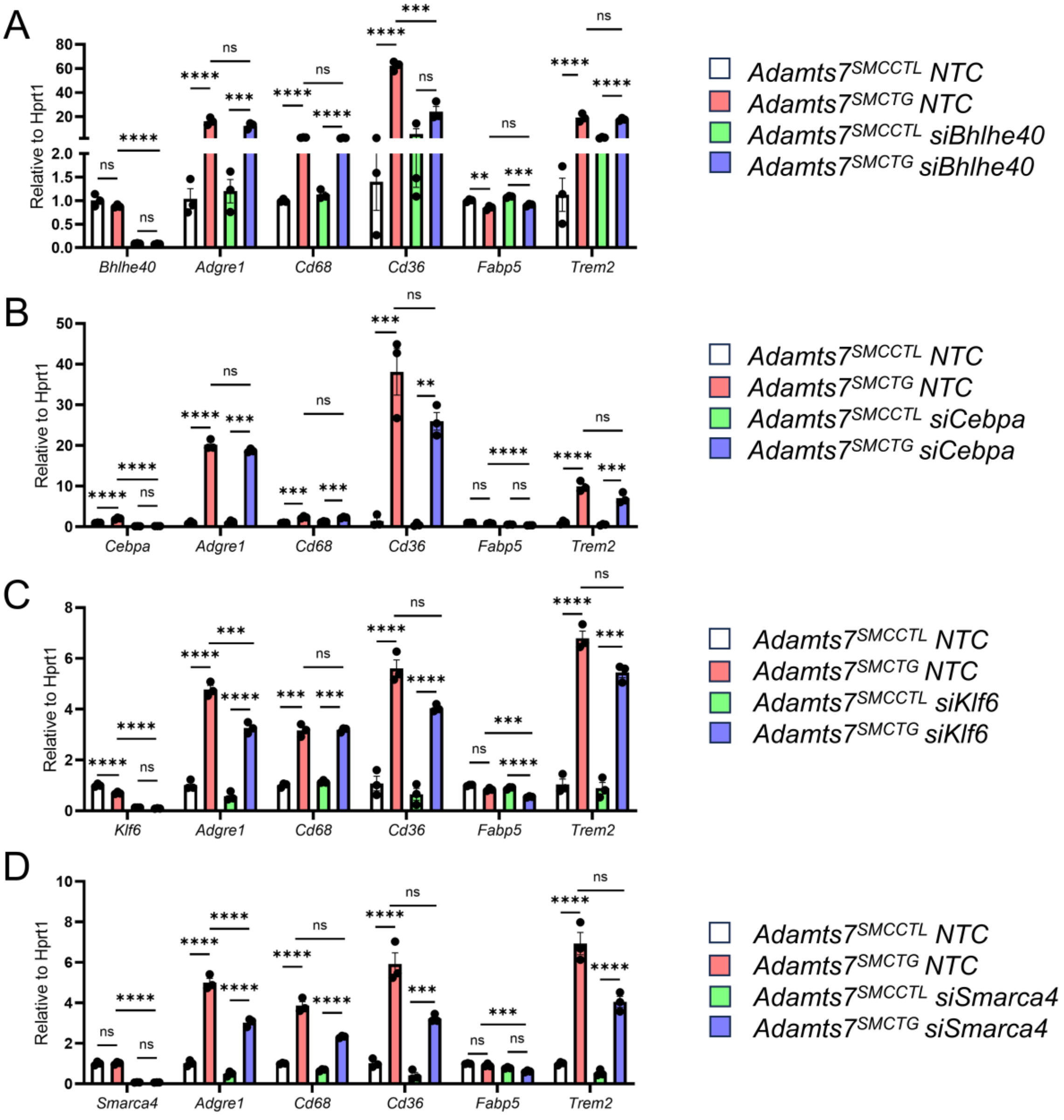


**Supplemental Figure 6. Knockdown of candidate transcription factors does not alter *Adamts7* conferred macrophage-like and lipid-handling genes.**

Indicated siRNAs (10nM) were used to knock down candidate transcription factors in primary SMCs from *Adamts7*$^{\text{SMCCTL}}$ and *Adamts7*$^{\text{SMCTG}}$ mice. 48 hours after transfection, cells were harvested for qRT-PCR analysis. ****$P$<0.0001, ***$P$<0.001, **$P$<0.01, *$P$<0.05 by two-way ANOVA with Bonferroni correction.

### SUPPLEMENTAL FIGURE 7

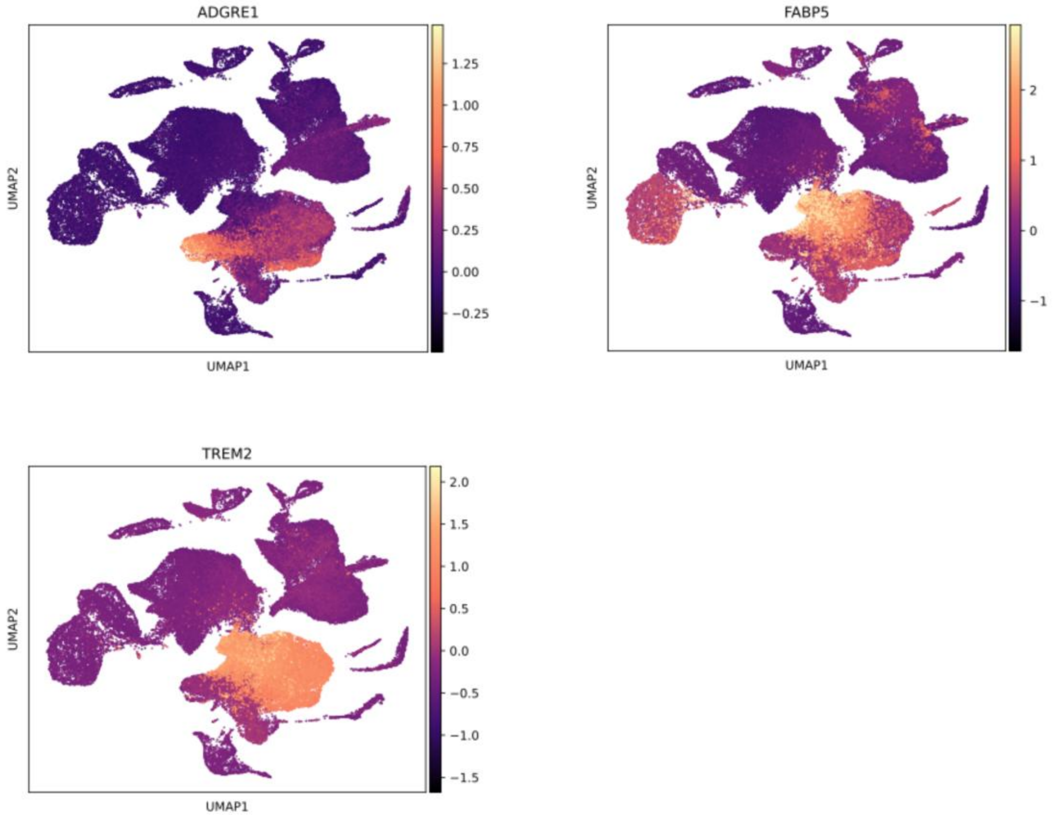


**Supplemental Figure 7. Lipid and macrophage-associated gene expression in human carotid atherosclerosis.**

Feature plots showing the expression of the indicated genes in scRNA-seq data from human carotid atherosclerotic plaques, as previously published by Bashore AC et al., Arterioscler Thromb Vasc Biol. 2024;44:930–945.

SUPPLEMENTAL FIGURE 8

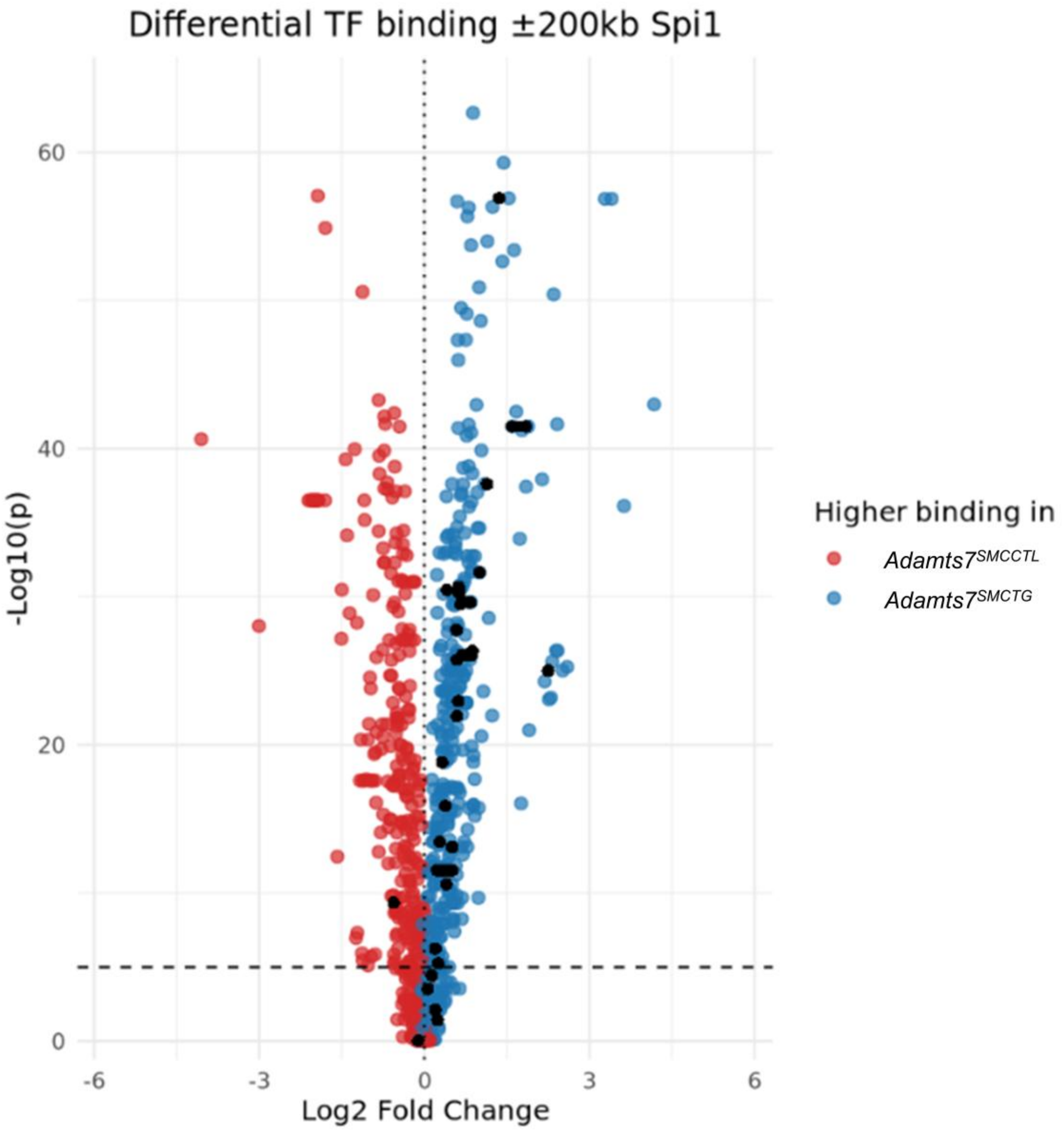


**Supplemental Figure 8. ATAC-seq reveals AP-1 binding motifs at the *Spi1* locus.**

Volcano plot showing differential chromatin accessibility in *Adamts7*$^{SMCCTL}$ and *Adamts7*$^{SMCTG}$ SMCs within ±200 kb of the *Spi1* transcription start site. Black dots indicate AP-1 binding motifs.

SUPPLEMENTAL FIGURE 9

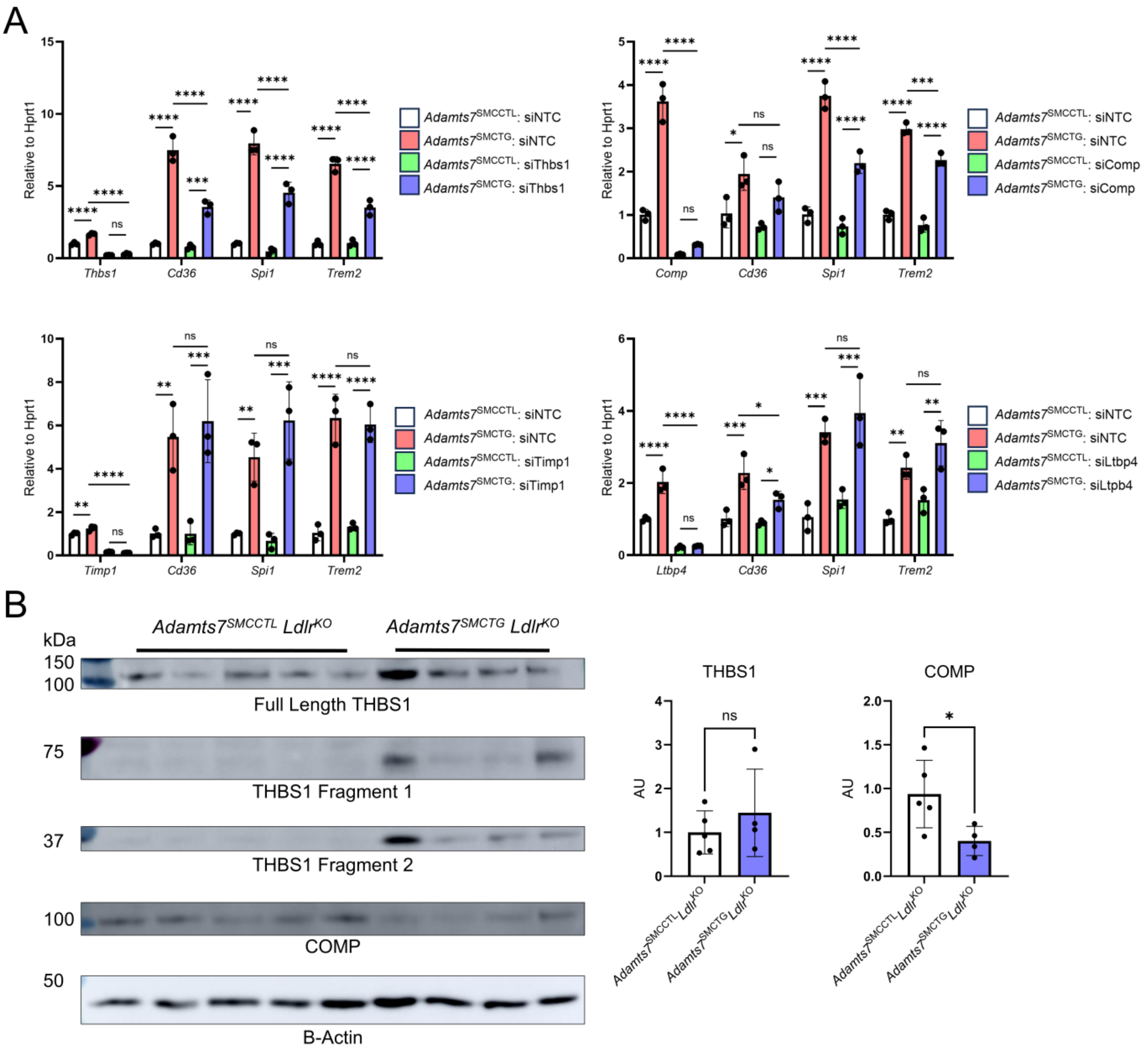

**Supplemental Figure 9. Knockdown of candidate ECM substrates to assess their role in SMCs.**

(A) Primary SMCs isolated from *Adamts7*$^{SMCCTL}$ and *Adamts7*$^{SMCTG}$ mice were transfected with the indicated siRNAs (10 nM). Cells were harvested 48 hours post-transfection for qRT-PCR analysis. Statistical significance was determined by two-way ANOVA with Sidak's multiple comparisons test: ****$P$<0.0001, ***$P$<0.001, **$P$<0.01, *$P$<0.05. (B) Western blot of whole-aorta lysates from mice fed a WTD, probed for ECM components THBS1 and COMP ($n$ = 4 - 5 male mice). Quantification of COMP and full-length THBS1 relative to B-actin. Statistical significance was assessed using a two-tailed Student's *t*-test. *$P$<0.05.
